## Supplementary material for "A multiscale model of complex endothelial cell dynamics in early angiogenesis": S1 Text

#### Contents

**1 Relating simulation units to experimental units** **1**

**2 Subcellular scale: VEGF-Delta-Notch signalling pathway** **2**

**3 Multiscale simulation algorithm** **9**

**4 Next Subvolume (NSV) method** **12**

**5 Metric definitions** **12**

**6 Mixing measure** **16**

**7 Quantification of simulated vascular networks** **22**

**8 Sensitivity analysis** **25**

### 1 Relating simulation units to experimental units

Based on [1, 2], we approximated the correspondence between time, space and VEGF concentration in our simulations and real experiments.

**Space** Since in our model we account for cell nucleus positions, we chose the voxel size such that it approximately corresponds to the size of the nucleus of an endothelial cell. Thus, we fixed the voxel width, $h = 5\ \mu m$.

**Time** In experiments from [1, 2], confocal microscopy imaging was carried out with time intervals of 15 minutes. Thus, in order to relate our simulation time to experimental time, we fix 0.03 of simulation time = 15 minutes. Then we calibrated our model, using this time correspondence, such that the dynamics of vascular growth in simulations and experiments from [1, 2] are in good quantitative agreement.

**VEGF concentration** In [1, 2], experiments were performed with uniform VEGF concentrations of 0, 5, and 50 ng/ml. Using molar mass of a VEGF molecule (40 kDa), the Avogadro constant ($N_A = 6.022 \times 10^{23}\ mol^{-1}$) and assuming that a cell is exposed to all VEGF molecules within a sphere of radius equal to 20 $\mu m$, simple computation yields the following correspondence

$$V = 0, \text{ external VEGF molecules} \sim 0\ ng/ml;$$
$$V = 2500, \text{ external VEGF molecules} \sim 5\ ng/ml;$$
$$V = 25000, \text{ external VEGF molecules} \sim 50\ ng/ml.$$

#### 2 Subcellular scale: VEGF-Delta-Notch signalling pathway

The subcellular scale of our model accounts for the phenotype specification mediated by the VEGF-Delta-Notch signalling pathway. Within a growing tumour, hypoxic regions of the tissue secrete tumour angiogenic factors (TAFs), which diffuse towards the underlying vascular bed and activate quiescent endothelial cells (ECs), thus inducing angiogenic sprouting [3, 4]. We focus on the vascular endothelial growth factor (VEGF), the most well-studied TAF. The canonical Delta-Notch pathway interacts with the VEGF pathway enabling a contact-mediated communication between ECs allowing them to coordinate their decision making processes [5, 6, 7, 8]. Specifically, ECs acquire one of the two phenotypes, tip or stalk, which are characterised by distinct gene expression patterns (tip cells: high Delta and VEGFR2, low Notch; stalk cell: low Delta and VEGFR2, high Notch) [5, 7, 9]. Phenotypic switching modifies EC behaviour which is manifested in cell-cell and cell-ECM interactions [3, 10]. In our model, these interactions are accounted for at the cellular and tissue scale. In turn, the local extracellular environment (configuration of the cellular and tissue scales) serves as a modulator of VEGF-Delta-Notch signalling [4, 11], thus, dynamical coupling between scales is maintained in both directions. In this section, we present a more detailed description of the formulation of the subcellular scale model.

Phenotype specification via the VEGF-Delta-Notch signalling pathway has been extensively investigated from the mathematical modelling perspective [12, 13, 14, 15, 16, 17, 18, 19, 20]. Here, we combine the lateral inhibition model of the Delta-Notch signalling pathway developed in [13, 14] with the VEGF signalling pathway as was done in [15, 16]. We refer to these works for a more detailed overview of the biological foundation of the model.

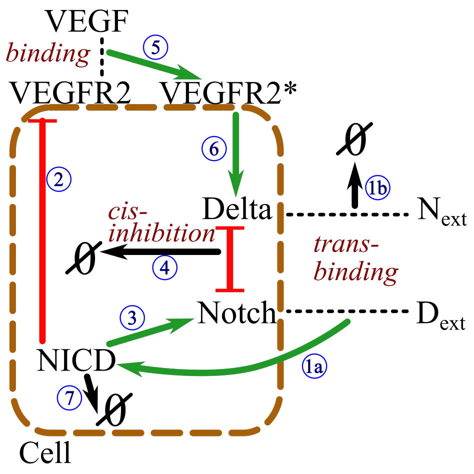


**SM-Fig 1. An illustration of the kinetic reactions of the VEGF-Delta-Notch signalling pathway for the individual cell system.** The reactions are labelled by blue circled numbers (see text for description). $N_{ext}$ and $D_{ext}$ represent extracellular levels of Notch and Delta, respectively.

##### Individual cell system

Let $N$ denote the level (number of proteins) of Notch receptor in a cell, $D$ – Delta ligand, $I$ – Notch intracellular domain (NICD), $R2$ – VEGF receptor 2 (VEGFR2) and $R2^*$ – activated, i.e. bound to VEGF, VEGFR2. We assume the focal cell is exposed to extracellular levels of Notch and Delta, $N_{ext}$ and $D_{ext}$, respectively (which belong to neighbouring cells, whose signalling dynamics we do not account for explicitly in the individual cell system, thus $N_{ext}$ and $D_{ext}$ are assumed to be constant). We denote by $V$ the extracellular level of VEGF. We assume $V$ to be constant as maintained in many *in vitro* experiments (e.g. [1, 2]). The kinetic reactions involved in our model are (see the illustration in SM-Fig 1):

(1a) $N + D_{ext} \underset{k_t^-}{\overset{k_t^+}{\rightleftarrows}} [N\,D_{ext}] \xrightarrow{k_p} I + D_{ext}$ — Trans-binding/unbinding of an extracellular Delta ligand to a Notch receptor on the focal cell and the following cleavage of a NICD.

(1b) $D + N_{ext} \underset{k_t^-}{\overset{k_t^+}{\rightleftarrows}} [D\,N_{ext}] \xrightarrow{k_p^1} D + N_{ext}$, $\;[D\,N_{ext}] \xrightarrow{k_p^2} N_{ext}$ — Trans-binding/unbinding of a Delta ligand on the focal cell to an extracellular Notch receptor on the focal cell followed by either endocytotic recycling of the Delta ligand (upper reaction with the rate $k_p^1$) or its degradation (lower reaction with the rate $k_p^2$).

(2) $\varnothing \underset{\gamma}{\overset{b_{R2} H^S(I;I_0,\lambda_{I,R2},n_{R2})}{\rightleftarrows}} R2$ — NICD-dependent production of a VEGFR2 receptor and its degradation at a constant rate.

(3) $\varnothing \underset{\gamma}{\overset{b_N H^S(I;I_0,\lambda_{I,N},n_N)}{\rightleftarrows}} N$ — NICD-dependent production of a Notch receptor and its degradation at a constant rate.

(4) $N + D \underset{k_c^-}{\overset{k_c^+}{\rightleftarrows}} [N\,D] \xrightarrow{k_e} \varnothing$ — Mutual cis-inhibition of a Delta ligand and a Notch receptor [13, 14].

(5) $R2 + V \xrightarrow{k_v} R2^* \xrightarrow{\gamma_e} \varnothing$ — Binding of an external VEGF to a VEGFR2 and the following degradation of the activated receptor.

(6) $\varnothing \underset{\gamma}{\overset{b_D H^S(R2^*;R2^*_0,\lambda_{R2^*,D},n_D)}{\rightleftarrows}} D$ — Production of a Delta ligand, dependent on activated (bound to VEGF) VEGFR2, and its degradation at a constant rate.

(7) $I \xrightarrow{\gamma_e} \varnothing$ — Degradation of a NICD.

Here $H^S(X; X_0, \lambda_{X,Y}, n_Y)$ is the so-called shifted Hill function

$$H^S(X; X_0, \lambda_{X,Y}, n_Y) = \frac{1 + \lambda_{X,Y}\left({}^{X}/_{X_0}\right)^{n_Y}}{1 + \left({}^{X}/_{X_0}\right)^{n_Y}}. \quad (1)$$

This function represents the transcriptional regulation of gene expression of a generic gene, $Y$, in response to the signalling variable $X$. $X_0$ and $n_Y$ are positive parameters characterising the activation threshold and cooperativity, respectively. For $0 \leqslant \lambda_{X,Y} < 1$ ($\lambda_{X,Y} > 1$) the production of $Y$ is down-regulated (up-regulated) as the amount of $X$ increases; if $\lambda_{X,Y} = 1$ then $X$ has no effect on production of $Y$ (see SM-Fig 2). We assume that NICD signalling up-regulates production of Notch, thus $\lambda_{I,N} > 1.0$ (reaction (3)), whereas production of VEGFR2 is down-regulated, $\lambda_{I,R2} < 1.0$ (reaction (2)). Activated VEGFR2 enhances Delta production, $\lambda_{R2*,D} > 1.0$ (reaction (6)). In these reactions, prefactors of the type $b_P$ characterise baseline production of the corresponding protein $P$.

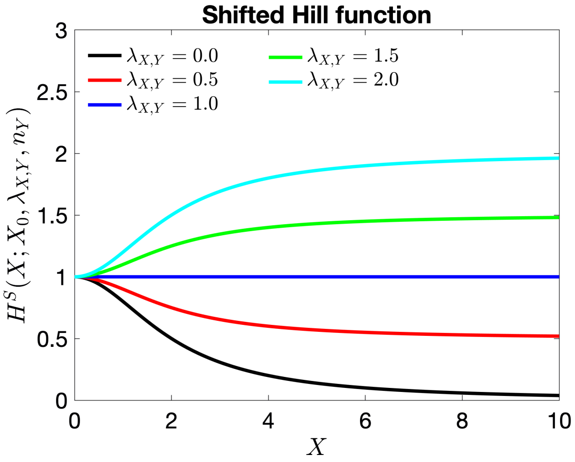


**SM-Fig 2. An illustration of the shifted Hill function for varying parameter $\lambda_{X,Y}$.** The functional form of $H^S(\cdot)$ is given by Eq (1). Here $X_0 = 2.0$, $n_Y = 2$.

We assume the same degradation rate, $\gamma$, for all inactivated proteins (Notch, Delta, VEGFR2). The half-life of activated signalling cues (NICD, activated VEGFR2) is much shorter, thus $\gamma_e > \gamma$ [16]. Furthermore, we assume that complexes [Delta-Notch] once formed release the signal or undergo endocytosis very fast compared to formation rate (i.e. $k_p^{1,2} \gg k_t^{\pm}$ and $k_e \gg k_c^{\pm}$). Thus, denoting $k_p = k_p^1 + k_p^2$, $k_t = k_t^+ \left(1 - \frac{k_t^-}{k_t^- + k_p}\right)$, $k_c = k_c^+ \left(1 - \frac{k_c^-}{k_c^- + k_e}\right)$ and $\eta = \frac{k_p^2}{k_p}$, we can simplify reactions (1a), (1b) and (4) :

(1a) $N + D_{ext} \xrightarrow{k_t} I + D_{ext}$

(1b) $D + N_{ext} \xrightarrow{(1-\eta)k_t} D + N_{ext}$, $\quad \searrow^{\eta k_t} N_{ext}$

(4) $N + D \xrightarrow{k_c} \varnothing$

To illustrate the characteristic behaviour of the individual cell system given by the kinetic reactions (1) -(7), we derive the associated mean-field limit equations:

$$\begin{aligned}
\frac{\mathrm{d}N}{\mathrm{d}t} &= b_N H^S(I; I_0, \lambda_{I,N}, n_N) - \gamma N - k_t D_{ext} N - k_c N D, \\
\frac{\mathrm{d}D}{\mathrm{d}t} &= b_D H^S(R2^*; R2^*_0, \lambda_{R2*,D}, n_D) - \gamma D - \eta k_t N_{ext} D - k_c N D, \\
\frac{\mathrm{d}I}{\mathrm{d}t} &= k_t D_{ext} N - \gamma_e I, \\
\frac{\mathrm{d}R2}{\mathrm{d}t} &= b_{R2} H^S(I; I_0, \lambda_{I,R2}, n_{R2}) - \gamma R2 - k_v V R2, \\
\frac{\mathrm{d}R2^*}{\mathrm{d}t} &= k_v V R2 - \gamma_e R2^*.
\end{aligned} \tag{2}$$

A full description and reference values of the parameters used in Eq (2) can be found in S1 Table.

To facilitate the analysis of the system of equations Eq (2), we introduce dimensionless variables and parameters (shown in SM-Table 1). The non-dimensional individual cell system reads

$$\begin{aligned}
\frac{\mathrm{d}n}{\mathrm{d}t} &= \beta_N H^S(\rho_N i; 1.0, \lambda_{I,N}, n_N) - n - d_{ext} n - \kappa n d, \\
\frac{\mathrm{d}d}{\mathrm{d}t} &= \beta_D H^S(\rho_{R2} r2^*; 1.0, \lambda_{R2*,D}, n_D) - d - \eta n_{ext} d - \kappa n d, \\
\frac{\mathrm{d}i}{\mathrm{d}t} &= d_{ext} n - \tau i, \\
\frac{\mathrm{d}r2}{\mathrm{d}t} &= \beta_{R2} H^S(\rho_N i; 1.0, \lambda_{I,R2}, n_{R2}) - (1 + v_{ext}) r2, \\
\frac{\mathrm{d}r2^*}{\mathrm{d}t} &= v_{ext} r2 - \tau r2^*.
\end{aligned} \tag{3}$$

For simplicity, we omit the bar in the dimensionless time variable.

We have studied numerically how the $d$- and $n$-nullclines of the system of equations Eq (3) change as we vary the (non-dimensional) external Delta concentration, $d_{ext}$. SM-Fig 3(A) shows that for low values of $d_{ext}$ (SM-Fig 3(A)(1)) lateral inhibition is not strong enough to suppress the default [6] tip phenotype. As

$d_{ext}$ increases, the system enters a bistable regime where both phenotypes, tip and stalk, coexist (see SM-Fig 3(A)(2)). Finally, for higher values of $d_{ext}$, the tip phenotype is suppressed and the only stable steady state of the system of equations Eq (3) corresponds to the stalk phenotype (see SM-Fig 3(A)(3)). A general picture can be seen in a numerically computed bifurcation diagram (SM-Fig 3(B)). The effect of external VEGF, $v_{ext}$, on phenotype specification is to enlarge the bistability region as its concentration increases (see SM-Fig 3(C)). On the contrary, for a fixed value of $v_{ext}$, increasing concentration of the external Notch, $n_{ext}$, reduces the size of the bistability region (see SM-Fig 3(D)).

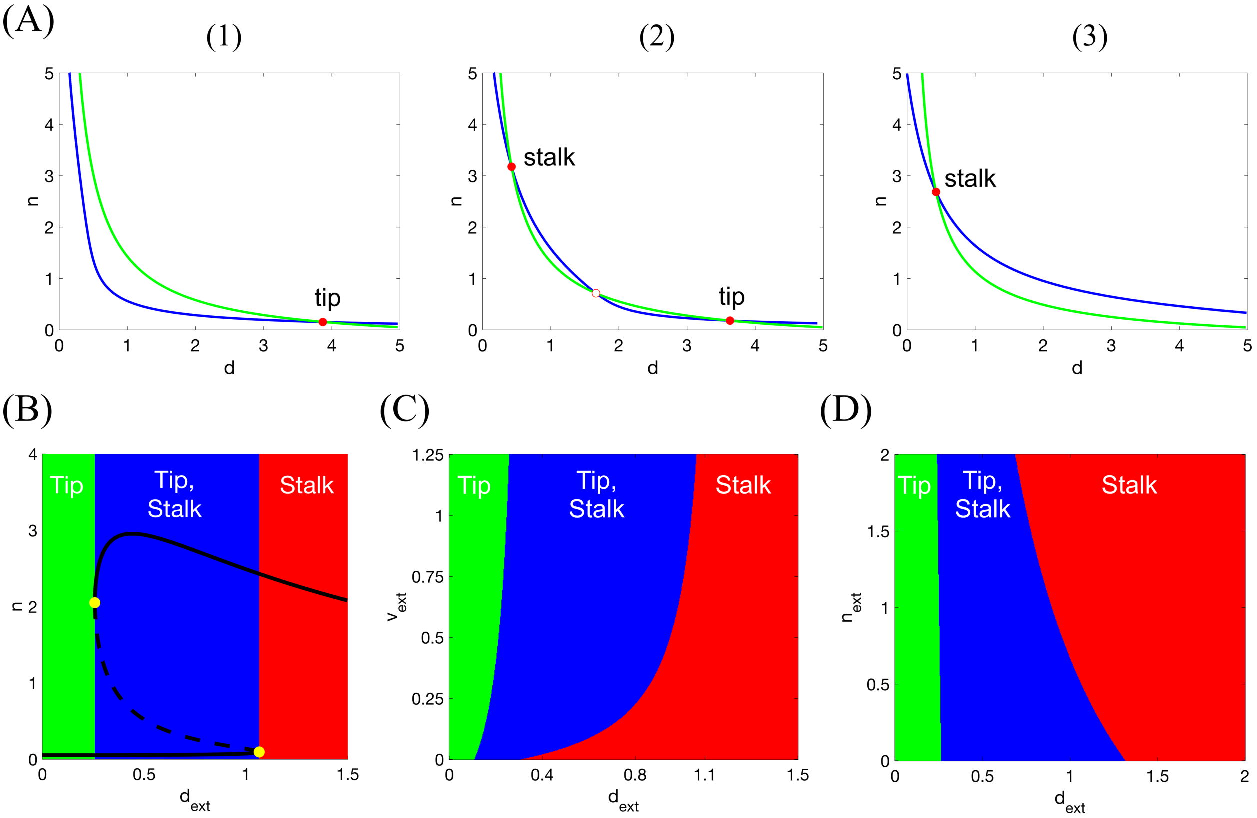


**SM-Fig 3. Numerical simulations of the non-dimensional mean-field single cell VEGF-Delta-Notch system (Eq (3)).** **(A)** $d$- and $n$-nullclines for varying level of external Delta, $d_{ext}$. **(1)** For low values of $d_{ext}$ (here $d_{ext} = 0.1$) there is only one (stable) steady state of the system corresponding to the tip phenotype. **(2)** A bistability region with two stable steady states: tip and stalk cells, occurs for intermediate values of $d_{ext}$ (here $d_{ext} = 0.3$). The unstable saddle point is indicated by a red unfilled circle. **(3)** For higher values of $d_{ext}$ (here $d_{ext} = 1.0$), the system is monostable with the stalk phenotype as its only (stable) steady state. **(B)** Bifurcation diagram of non-dimensional Notch concentration, $n$, as a function of external Delta ligand, $d_{ext}$, corresponding to the system of equations Eq (3). Full lines denote stable steady states; dashed lines – unstable steady state; yellow filled dots – saddle-node bifurcation points. **(C)** Phenotype diagram as a function of external Delta, $d_{ext}$, and external VEGF, $v_{ext}$, corresponding to the system of equations Eq (3). **(D)** Phenotype diagram as a function of external Delta, $d_{ext}$, and external Notch, $n_{ext}$, corresponding to the system of equations Eq (3). Parameter values used to make the plots (except for those indicated specifically) are listed in SM-Table 1.

| **Variable/ Parameter** | **Ref. value** | **Description** |
| --- | --- | --- |
| $N_0 = \frac{\gamma}{k_t}$ | 2000.0 | The characteristic level of Notch. |
| $D_0 = \frac{\gamma}{k_t}$ | 2000.0 | The characteristic level of Delta. |
| $R2_0 = \frac{\gamma}{k_v}$ | 2000.0 | The characteristic level of VEGFR2. |
| $\rho_N = \frac{N_0}{I_0}$ | 20.0 | The ratio of the characteristic levels of Notch and NICD. |
| $\rho_{R2} = \frac{R2_0}{aR2_0}$ | 10.0 | The ratio of the characteristic levels of unbound and bound VEGFR2. |
| $n = \frac{N}{N_0}$ | | Non-dimensional Notch receptor concentration. |
| $D = \frac{D}{D_0}$ | | Non-dimensional Delta ligand concentration. |
| $i = \frac{I}{N_0}$ | | Non-dimensional NICD concentration. |
| $r2 = \frac{R2}{R2_0}$ | | Non-dimensional VEGFR2 concentration. |
| $r2^* = \frac{R2^*}{R2_0}$ | | Non-dimensional VEGF-bound VEGFR2 concentration. |
| $\beta_N = \frac{b_N k_t}{\gamma^2}$ | 2.5 | Non-dimensional baseline expression of Notch receptor. |
| $\beta_D = \frac{b_D k_t}{\gamma^2}$ | 4.0 | Non-dimensional baseline expression of Delta ligand. |
| $\beta_{R2} = \frac{b_N k_v}{\gamma^2}$ | 4.0 | Non-dimensional baseline expression of VEGFR2. |
| $v_{ext} = \frac{k_v}{\gamma} V = \frac{V}{R2_0}$ | 1.25 | Non-dimensional external VEGF concentration. |
| $d_{ext} = \frac{D_{ext}}{D_0}$ | 0.0 – 5.0 | Non-dimensional external Delta ligand concentration. |
| $n_{ext} = \frac{N_{ext}}{N_0}$ | 0.0 – 5.0 | Non-dimensional external Notch receptor concentration. |
| $\kappa = \frac{k_c}{k_t}$ | 12.0 | The ratio of cis- and trans-binding for Delta and Notch. According to [13], $\kappa > 1$. |
| $\tau = \frac{\gamma_e}{\gamma}$ | 5.0 | The ratio of degradation rates of activated and non-activated receptors/ligands. Activated signals always have shorter half-life, thus $\tau > 1$. |
| $\bar{t} = \gamma t$ | | Non-dimensional time. |

**SM-Table 1. Non-dimensional variables and parameters of the VEGF-Delta-Notch system.** The reference values are computed according to the dimensional parameter values listed in S1 Table. For the multicellular system external Notch and Delta, $n_{ext}$ and $d_{ext}$, respectively, vary according to the configuration of the system (see text for details).

#### Multicellular system

The individual cell model can be easily extended to a multicellular one. We need to specify the external levels of Delta and Notch ($D_{ext}$ and $N_{ext}$, respectively) to which each individual cell is exposed. $D_{ext}$ and $N_{ext}$ are given by the levels of the corresponding proteins summed over all the neighbouring cells with which the focal cell is in contact. Since in our multiscale model of angiogenesis we account for cell nucleus positions instead of the exact cell membrane configurations, cell-cell interactions are assumed to be non-local, i.e. beyond their first neighbours. We define an interaction radius, $R_s$, and assume that two cells are in contact if they are contained (totally or partially) within a circular neighbourhood of radius $R_s$. Taking into account that we use an on-lattice modelling approach with a uniform hexagonal discretisation of the domain, we define a local neighbourhood of a cell located at the voxel with an index $k \in \mathbb{R}^2$, $v_k$, as a set of voxels

$$H(k) := \{v_l : v_l \cap \mathcal{B}_{R_s}(k) \neq \varnothing,\ l \neq k,\ l \in \mathcal{I}\}, \tag{4}$$

where $\mathcal{B}_{R_s}(k)$ denotes a circular neighbourhood of radius $R_s$ centred at the centre of voxel $k$, and $\mathcal{I}$ is the set of all voxel indices.

The amount of Delta on a neighbour $l \in H(k)$ which is in contact with Notch receptors of the cell of interest $k$ is assumed to be proportional to the surface area of the overlap between the circular neighbourhood of the focal cell $k$ and the neighbour $l$. This is given by the weights $\alpha_{kl}$ (see SM-Fig 4) defined as follows

$$\alpha_{kl} = \frac{|v_l \cap \mathcal{B}_{R_s}(k)|}{|v_l|}, \quad k,\ l \in \mathcal{I}. \tag{5}$$

Here $|\cdot|$ stands for the surface area.

Thus, the external Delta (Notch) concentration, $D_{ext}$ ($N_{ext}$), for a cell situated in a voxel $v_k$ is defined as follows

$$\begin{aligned} D_{ext} &= \overline{D_k} = \sum_{l\ \in\ H(k)} \alpha_{kl} D_l, \\ N_{ext} &= \overline{N_k} = \sum_{l\ \in\ H(k)} \alpha_{kl} N_l. \end{aligned} \tag{6}$$

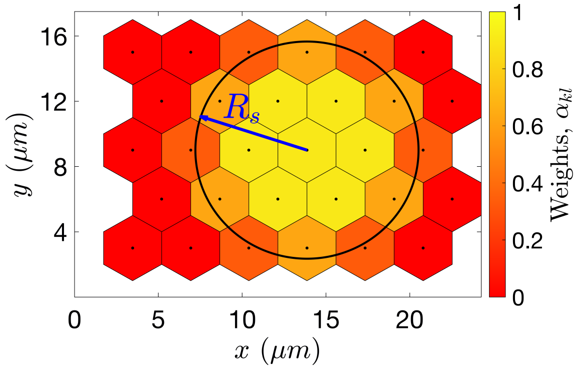


**SM-Fig 4. Illustration of values of the weight coefficients, $\boldsymbol{\alpha_{kl}}$ (see Eq (5)).** $R_s$ is the interaction radius. The colour bar indicates the value of the weight, $\alpha_{kl}$, for each of the neighbours in the lattice.

We can now rewrite the kinetic reactions ①-⑦ for the multicellular system in a straightforward manner. For each cell in the system, positioned in voxel $v_k$, we consider the following kinetic reactions (numbered with Roman numerals as equivalents of the kinetic reactions of the individual cell system numbered with Arabic numerals)

(I) $N_k + D_l \xrightarrow{\alpha_{kl}(1-\eta)k_t} I_k + D_l$ $\qquad l \in H(k)$

$\qquad \searrow^{\alpha_{kl}\eta k_t}\ I_k$

(II) $\varnothing \underset{\gamma}{\overset{b_{R2}H^S(I_k; I_0, \lambda_{I,R2}, n_{R2})}{\rightleftarrows}} R2_k$

(III) $\varnothing \underset{\gamma}{\overset{b_N H^S(I_k; I_0, \lambda_{I,N}, n_N)}{\rightleftarrows}} N_k$

(IV) $N_k + D_k \xrightarrow{k_c} \varnothing$

(V) $R2_k + V \xrightarrow{k_v} R2^*_k \xrightarrow{\gamma_e} \varnothing$

(VI) $\varnothing \underset{\gamma}{\overset{b_D H^S(R2^*_k; R2^*_0, \lambda_{R2^*,D}, n_D)}{\rightleftarrows}} D_k$

(VII) $I_k \xrightarrow{\gamma_e} \varnothing$

Note that now the reactions (1a) and (1b) of the individual cell system result in the same reaction (I) of the multicellular model. Reaction (I) is bimolecular between a Notch receptor in voxel $v_k$ and a Delta ligand in a neighbouring cell in $v_l$. As a result of this reaction, a NICD is produced in the cell to which the Notch receptor belongs (voxel $v_k$) and Delta on the neighbour, $D_l$, is either degraded or endocytotically recycled.

To summarise, the subcellular scale of our model is a stochastic system given by the multicellular system with kinetic reactions (I)-(VII). We simulate it using a variation of the Stochastic Simulation Algorithm (SSA), the Next Subvolume method (NSV) [21]. We list transition rates and the corresponding stoichiometric vectors used for simulation in SM-Table 2.

When simulated in a simple two-dimensional domain, with only first-neighbour (voxels that share an edge) interactions, the multicellular system (reactions (I)-(VII)) produces the classical chessboard pattern [12] of alternating tip/stalk cells. However, our system amplifies the range of possible patterns beyond the classical one. This is due to accounting for mutual cis-inhibition of Delta and Notch within the same cell and non-locality of interactions within the radius $R_s$. Specifically, increasing the cis-binding parameter, $k_c$, allows tip cells to be neighbours since increasing $k_c$ reduces the lateral inhibition ability of cells (see SM-Fig 5). In contrast, increasing interaction radius, $R_s$, enhances the ability of a tip cell to inhibit more neighbours and prevent them from acquiring the tip phenotype. Thus, the distance between two tip cells increases as $R_s$ grows (see S1 Fig).

For completeness, we list the mean-field limit equations corresponding to the multicellular kinetic reactions (I)-(VII):

$$\begin{aligned}
\frac{\mathrm{d}N_k}{\mathrm{d}t} &= b_N H^S(I_k; I_0, \lambda_{I,N}, n_N) - \gamma N_k - k_t \overline{D_k} N_k - k_c N_k D_k,\\
\frac{\mathrm{d}D_k}{\mathrm{d}t} &= b_D H^S(R2^*_k; R2^*_0, \lambda_{R2*,D}, n_D) - \gamma D_k - \eta k_t \overline{N_k} D_k - k_c N_k D_k,\\
\frac{\mathrm{d}I_k}{\mathrm{d}t} &= k_t \overline{D_k} N_k - \gamma_e I_k,\\
\frac{\mathrm{d}R2_k}{\mathrm{d}t} &= b_{R2} H^S(I_k; I_0, \lambda_{I,R2}, n_{R2}) - \gamma R2_k - k_v V R2_k,\\
\frac{\mathrm{d}R2^*_k}{\mathrm{d}t} &= k_v V R2_k - \gamma_e R2^*_k.
\end{aligned} \tag{7}$$

Here $\overline{D_k}$ and $\overline{N_k}$ are given by Eq (6) and $k \in \mathcal{I}$.

| Reaction, $\boldsymbol{R}$ | Transition rate | $\boldsymbol{\nu_k^R}$ | $\boldsymbol{\nu_l^R}$ |
|---|---|---|---|
| I | $\frac{(1-\eta)\alpha_{kl}k_t}{\Omega}N_k D_l$ | $(-1,0,+1,0,0)^T$ | $(0,0,0,0,0)^T$ |
| | $\frac{\eta\alpha_{kl}k_t}{\Omega}N_k D_l$ | $(-1,0,+1,0,0)^T$ | $(0,-1,0,0,0)^T$ |
| II | $\Omega b_{R2}H^S(I_k;\Omega I_0,\lambda_{I,R2},n_{R2})$ | $(0,0,0,+1,0)^T$ | $(0,0,0,0,0)^T$ |
| | $\gamma R2_k$ | $(0,0,0,-1,0)^T$ | $(0,0,0,0,0)^T$ |
| III | $\Omega b_N H^S(I_k;\Omega I_0,\lambda_{I,N},n_N)$ | $(+1,0,0,0,0)^T$ | $(0,0,0,0,0)^T$ |
| | $\gamma N_k$ | $(-1,0,0,0,0)^T$ | $(0,0,0,0,0)^T$ |
| IV | $\frac{k_c}{\Omega}N_k D_k$ | $(-1,-1,0,0,0)^T$ | $(0,0,0,0,0)^T$ |
| V | $VR2_k$ | $(0,0,0,-1,+1)^T$ | $(0,0,0,0,0)^T$ |
| | $\gamma_e R2_k^*$ | $(0,0,0,0,-1)^T$ | $(0,0,0,0,0)^T$ |
| VI | $\Omega b_D H^S(R2_k^*;\Omega R2_0^*,\lambda_{R2*,D},n_D)$ | $(0,+1,0,0,0)^T$ | $(0,0,0,0,0)^T$ |
| | $\gamma D_k$ | $(0,-1,0,0,0)^T$ | $(0,0,0,0,0)^T$ |
| VII | $\gamma_e I_k$ | $(0,0,-1,0,0)^T$ | $(0,0,0,0,0)^T$ |

**SM-Table 2. Transition rates of the multicellular VEGF-Delta-Notch system.** The transition rates are appropriately scaled with the system size parameter, $\Omega$ (in our simulations, we fix $\Omega = 200$). $\nu_r^R$ denotes a stoichiometric vector corresponding to a reaction $R$ in a cell at voxel $v_r$ indexed as $(N, D, I, R2, R2^*)^T$. The transition rates are calculated for all $k \in \mathcal{I}$ and $l \in H(k)$, where $\mathcal{I}$ denotes the set of voxel indices in the system.

### 3 Multiscale simulation algorithm

For the formulation of the algorithm we use the following notation:

| | |
|---|---|
| $\mathcal{I}$ | The set of voxel indices in the lattice, $\mathcal{L}$. |
| $N_I$ | The total number of voxels in the lattice, $\mathcal{L}$. |
| $\mathcal{C}^s = (\boldsymbol{N}, \boldsymbol{D}, \boldsymbol{I}, \boldsymbol{R2}, \boldsymbol{R2^*})$ | The full-lattice configuration of the variables of the subcellular scale: Notch, $\boldsymbol{N} = (N_1, \ldots, N_{N_I})$; Delta, $\boldsymbol{D} = (D_1, \ldots, D_{N_I})$; NICD, $\boldsymbol{I} = (I_1, \ldots, I_{N_I})$; VEGFR2, $\boldsymbol{R2} = (R2_1, \ldots, R2_{N_I})$; activated (bound to VEGF) VEGFR2, $\boldsymbol{R2^*} = \left(R2_1^*, \ldots, R2_{N_I}^*\right)$. |
| $\mathcal{C}_i^s = (N_i, D_i, I_i, R2_i, R2_i^*)$ | The configuration of the variables of the subcellular scale in voxel $v_i$. |
| $\mathcal{C}^c = (\boldsymbol{E}, \boldsymbol{D}, \boldsymbol{c}, \boldsymbol{m}, \boldsymbol{l})$ | The full-lattice configuration of the following variables: cellular scale cell distribution, $\boldsymbol{E} = (E_1, \ldots, E_{N_I})$; subcellular Delta, $\boldsymbol{D} = (D_1, \ldots, D_{N_I})$, used as a proxy to define cell phenotype; tissue scale ECM concentration, $\boldsymbol{c} = (c_1, \ldots, c_{N_I})$; tissue scale BM component concentrations, $\boldsymbol{m} = (m_1, \ldots, m_{N_I})$; tissue scale orientation landscape variable of ECM fibril alignment, $\boldsymbol{l} = (l_1, \ldots, l_{N_I})$. |
| $\omega(i \to j)$ | Transition rate for a migration event from voxel $v_i$ to voxel $v_j$. |
| $\tau_{ij}$ | Time step corresponding to the transition $\omega(i \to j)$. |

| | |
|---|---|
| $\tau$ | Time step of the next transition to occur. |
| $\text{swap}(A_i, A_j)$ | A swap operator for the variable $A$ exchanging its values corresponding to voxels $v_i$ and $v_j$, respectively. |
| $t$ | Simulation time. |
| $T_{max}$ | Final simulation time. |
| $N_R$ | Number of realisations. |

**Algorithm 1.** Pseudocode algorithm of multiscale model simulations.

1: Specify simulation setup.
2: Nullify the realisation counter, $count_R = 0$.
3: **while** $count_R < N_R$ **do**
4: $count_R = count_R + 1$.
5: Initialisation.
6: Set $t = 0$, $\tau = 1.0$.
7: **while** $t < T_{max}$ **do**
8: Obtain $\mathcal{C}^s$ by simulating the subcellular VEGF-Delta-Notch multicellular system with the final simulation time, $\tau$.
9: Given $\mathcal{C}^c$, compute migration transition rates, $\omega(i \to j)$, (see Eq (7) of the main text) for all $i, j \in \mathcal{I}$.
10: Sample waiting times for each transition from the Poisson distribution with the intensity given by the corresponding transition rate, $\tau_{ij} = Poisson(\omega(i \to j))$, for all $i, j \in \mathcal{I}$ such that $\omega(i \to j) > 0$.
11: Find the migration event with the minimum waiting time: $\tau = \tau_{\bar{i}\bar{j}} = \min_{i,j} \tau_{ij}$.
Set the jump direction vector, $s = h^{-1}(q_{\bar{j}} - q_{\bar{i}})$, where $h$ is the voxel width.
12: Perform the migration event: $\text{swap}(E_{\bar{i}}, E_{\bar{j}})$ and $\text{swap}(\mathcal{C}^s_{\bar{i}}, \mathcal{C}^s_{\bar{j}})$.
13: Update the orientation landscape due to traction forces generated by the migration event according to Eq (16) (of the main text) with $i = \bar{i}$ and $j = \bar{j}$ and the migration direction $s$.
14: Do a general update of the tissue scale variables with the final time $\tau$: fibrils relaxation, $\boldsymbol{l}$, (Eq (17) of the main text); ECM concentration, $\boldsymbol{c}$, (Eq (18)-(19) of the main text); BM component concentration, $\boldsymbol{m}$, (Eq (20)-(21) of the main text).
15: Increment the simulation time: $t = t + \tau$.
16: [Optional] Calculate statistics.
17: **end while**
18: [Optional] Post-processing and statistical analysis.
19: End of the current realization.
20: **end while**
21: End of simulation.

For clarity, we add a few comments to the pseudocode Algorithm 1.

*line 1* Simulation setup is defined by specifying the lattice dimensions, $N_I^x$ and $N_I^y$, the set of indices corresponding to the vascular plexus, $\mathcal{I}_{VP}$, the initial cell nucleus positions, $\mathcal{I}_{init}$, the initial ECM alignment, $s_{init}$, the initial ECM and BM component concentrations, $c_{init}$ and $m_{init}$, respectively, the distribution of VEGF, $V$, and the final simulation time, $T_{max}$. In our simulations, we used 4 different simulation setups, as listed in S4 Table.

*line 5* Initialisation of all variables is done as specified in S3 Table.

*line 6* In this line we set $\tau = 1.0$. This is used as the final simulation time for the first simulation of the subcellular VEGF-Delta-Notch system, i.e. for the initial phenotype patterning. Initial pattern

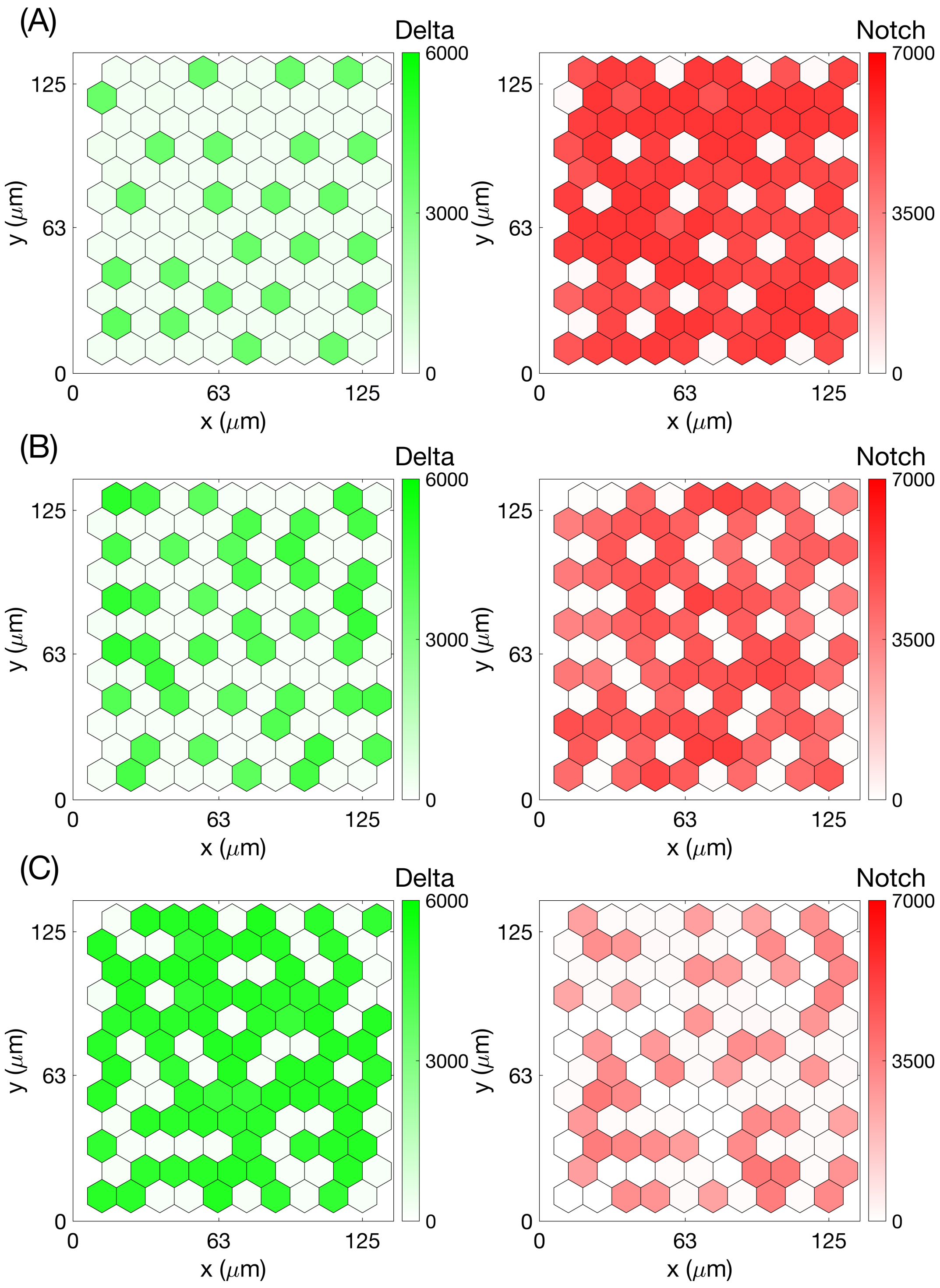


**SM-Fig 5. Examples of steady state patterns of the VEGF-Delta-Notch multicellular system for different cis-inhibition parameter values.** Final steady state patterns established during single stochastic simulations of the system described by the kinetic reactions (I)-(VII) for a uniform hexagonal lattice of $10 \times 12$ voxels. Cis-inhibition parameter was taken as **(A)** $k_c = 6.0e-4$; **(B)** $k_c = 10.0e-4$; **(C)** $k_c = 15.0e-4$. External VEGF level, $V = 2500$, and the rest of the parameter values as in S1 Table.

stabilisation from a uniform configuration set in the initialisation takes longer than when a pre-pattern exists and only several cells change their positions. Thus a value of 1.0 was chosen as a value sufficiently larger than a typical time step for a migration event (defined in *line 8*).

*line 8* The subcellular VEGF-Delta-Notch system is simulated using the Next Subvolume (NSV) method with the final time given by the time step of the last migration transition, $\tau$ (except for the first iteration step in which we use $\tau = 1.0$).

*line 10* An equivalent procedure to sample a number, $x$, from the Poisson distribution with intensity $\lambda$ is $x = {}^{1}/_{\lambda} \log \left( {}^{1}/_{\mathrm{Unif}[0,1]} \right)$, where Unif$[0,1]$ is the uniform distribution on $[0,1]$.

*line 12* The swapping events account for both a simple migration event, when the destination voxel $v_{\bar{j}}$ is free and a cell from voxel $v_{\bar{i}}$ is just put in a new position (voxel $v_{\bar{j}}$), and a switching event, when the destination voxel $v_{\bar{j}}$ is occupied and cells exchange their positions (the values of the subcellular scale).

### 4 Next Subvolume (NSV) method

The Next Subvolume (NSV) method is one of the modifications of the standard Stochastic Simulation Algorithm (SSA) [22], introduced in [21]. For reaction-diffusion systems, it is more efficient than the SSA since the computational effort grows as the logarithm of the number of voxels (subvolumes in 3D) instead of the linear dependency exhibited by the SSA.

In the SSA the time step for the next reaction to occur is generated from the Poisson distribution with the intensity equal to the total propensity for all voxels in the system. Then the reaction is chosen probabilistically according to the weight given by the local (for each voxel separately) propensities of each reaction. A major advantage of the NSV method is that time steps for the next reaction are generated for each voxel separately and stored in a sorted way according to the next time for a reaction to occur. Implementation of this algorithm is usually done by utilising a data structure called “priority queue” for which many efficient algorithms exist. This decreases the overall complexity of the simulating algorithm from linear (for the SSA) to logarithmic (for the NSV method) of the total number of voxels.

### 5 Metric definitions

Here we provide more details on definitions of and computational algorithms for the metrics used for model calibration. Firstly, we recall some of the notation used in the main text.

| | |
|---|---|
| $\mathcal{I}$ | Total set of voxel indices. |
| $v_i$ | A voxel indexed by $i \in \mathcal{I}$. |
| $\boldsymbol{E} = (E_1, \ldots, E_{N_I})$ | Distribution of cell nuclei. Here $N_I$ denotes cardinal of $\mathcal{I}$. |
| $T_{max}$ | Final simulation time in a single realization of our model. |
| $\iota$ | A cell label. |

We introduce a partitioning of a simulation time interval $[0, T_{max}]$ with a uniform step $\Delta$ as follows:

$$\mathcal{T}(\Delta) = \left\{ t_k = \Delta k,\ k = 0, \ldots, K,\ K = \left\lfloor \frac{T_{max}}{\Delta} \right\rfloor \right\}, \tag{8}$$

where $\lfloor x \rfloor$ denotes the largest integer less than or equal to $x \in \mathbb{R}$.

The pseudocode algorithms shown below should be considered as complementary to the general algorithm of multiscale simulations, Algorithm 1.

We also note that since the simulations we perform are stochastic, the statistic corresponding to $t_k \in \mathcal{T}(\Delta)$ is calculated at time $\bar{t}$ such that $t < t_k < \bar{t}$, where $t$ and $\bar{t}$ are time moments corresponding to two consecutive times of migration events at the cellular scale of the multiscale simulation.

**Displacement** The displacement statistic characterises the average displacement a cell makes in $\Delta_{disp}$ time. In our simulations, $\Delta_{disp}$ is taken such that it corresponds to 15 minutes, in order to be able to compare to the data in [2]. However, in general, $\Delta_{disp}$ can be chosen arbitrary.

The general algorithm to compute this statistic is given in Algorithm 2. Therein, the concatenation operator, $\cdot \vee \cdot$, is defined as $v^1 \vee v^2 = \left(v_1^1, \dots v_{N_1}^1, v_1^2, \dots v_{N_2}^2\right)$ for vectors $v^1 = \left(v_1^1, \dots v_{N_1}^1\right)$ and $v^2 = \left(v_1^2, \dots v_{N_1}^2\right)$.

**Algorithm 2.** Pseudocode algorithm for computing the displacement statistic.

Specify the length of the displacement interval, $\Delta_{disp}$.
Create an empty vector of displacements, $V_{disp} = \varnothing$.
**for** each realization **do**
  Nullify the simulation time, $t = 0$.
  $k = 1$.
  $V^1 = (0, \dots, 0) \in \mathbb{R}^{max\_cell}$, where $max\_cell = \sum_{i \in \mathcal{I}} E_i(t)$.
  **while** $t < T_{max}$ **do**
    Obtain time step, $\tau$, for the next migration event, $\omega(i \to j)$, (see Algorithm 1).
    Identify labels of cells whose nuclei are in voxels $v_i$ and $v_j$, $\iota_i$ and $\iota_j$, respectively, (if $E_j(t) = 0$, i.e. $v_j$ is empty, then only $\iota_i$).
    Increment the components of the displacement vector, corresponding to the labels of migrating cells, $V_{\iota_i}^k = V_{\iota_i}^k + h$ and $V_{\iota_j}^k = V_{\iota_j}^k + h$, where $h$ is the voxel width (if $E_j(t) = 0$, i.e. $v_j$ is empty, then only increment $V_{\iota_i}^k$).
    $t = t + \tau$.
    **if** $t > \Delta_{disp} k$ **then**
      $V_{disp} = V_{disp} \vee V^k$.
      $k = k + 1$.
      $V^k = (0, \dots, 0) \in \mathbb{R}^{max\_cell}$, where $max\_cell = \sum_{i \in \mathcal{I}} E_i(t)$.
    **end if**
  **end while**
**end for**
The output vector, $V_{disp}$, is a sample of cell displacements during all time intervals of length $\Delta_{disp}$ for all cells in all realisations. We use it to compute a probability density function of displacement in $\Delta_{disp}$ time (as in Fig 10(A) of the main text).

**Orientation** This statistic is used to characterise persistence of cell migration. The following procedure is used for its computation.

Each individual cell in a single realization is associated with a label, $\iota$. We record its trajectory within the lattice during the simulation, $p(\iota, t) \in \mathbb{R}^2$, where $t \in [0, T_{max}]$.

Each $p(\iota, t)$ (or any sample extracted from it) is a polygonal chain, since cells perform jumps at discrete time moments and are assumed to be motionless between them. We define the length of a polygonal chain $p(\iota, t)$ given the time partitioning $\mathcal{T}_\iota$ as follows

$$l(p(\iota, t) \mid \mathcal{T}_\iota) = \sum_k \|p(\iota, t_{k+1}) - p(\iota, t_k)\|, \qquad t_k \in \mathcal{T}_\iota.$$

Here $\|\cdot\|$ is the Euclidean norm in $\mathbb{R}^2$.

We denote by $\mathcal{T}_\iota^r = \{t_k\}_k$ the ascending sequence of time moments of migration events of the cell with label $\iota$. Thus, $\mathcal{T}_\iota^r$ defines the real trajectory of the cell $\iota$ (see SM-Fig 6(A)).

We also define a partitioning of the simulation time interval, defining the smoothed trajectory (see SM-Fig 6(B)), as $\mathcal{T}_\iota^s = \mathcal{T}(\Delta_{orient})$ (using Eq (8)), where $\Delta_{orient}$ is a uniform time step. In order to be able to compare our simulation results with experimental data from [1], $\Delta_{orient}$ was chosen such that it corresponds to 20 minutes. However, in general, $\Delta_{orient}$ can be arbitrary provided that it is greater than a typical waiting time between migration events in our multiscale simulation algorithm.

Then the orientation quantity, $O_\iota$, is defined as

$$O_\iota = \frac{l(p(\iota, t) \mid \mathcal{T}_\iota^s)}{l(p(\iota, t) \mid \mathcal{T}_\iota^r)}.$$

When $O_\iota$ is close to 1.0, the cell $\iota$ is characterised as persistent. Lower values of $O_\iota$ correspond to trajectories in which cells performed many backward jumps (in the direction opposite to the elongation direction of a sprout).

The quantities $O_\iota$ are computed for each individual cell in each realization. The overall sample of these quantities is used to produce box plots of the type shown in Fig 10(C) of the main text.

**Directionality** The directionality statistic provides a breakdown of migration events by their direction with respect to the direction of sprout elongation. We account for three types of movement: anterograde, retrograde and no movement (still). Since in our simulations ECs polarise according to the alignment of ECM fibrils (orientation landscape variable) which defines the direction of sprout elongation, any movement of a cell from the voxel of origin is considered anterograde. In contrast, if there is a cell in the voxel of destination of the migration event, then this cell is overtaken and is displaced to the voxel of origin. Thus, this is a retrograde movement. To define cells that stay still, we introduce a parameter, $\Delta_{still}$. A cell is assumed to be still if it has not moved in $\Delta_{still}$ time (in our simulations this parameter corresponds to 20 minutes). A pseudocode for computing the directionality statistic is shown in Algorithm 3.

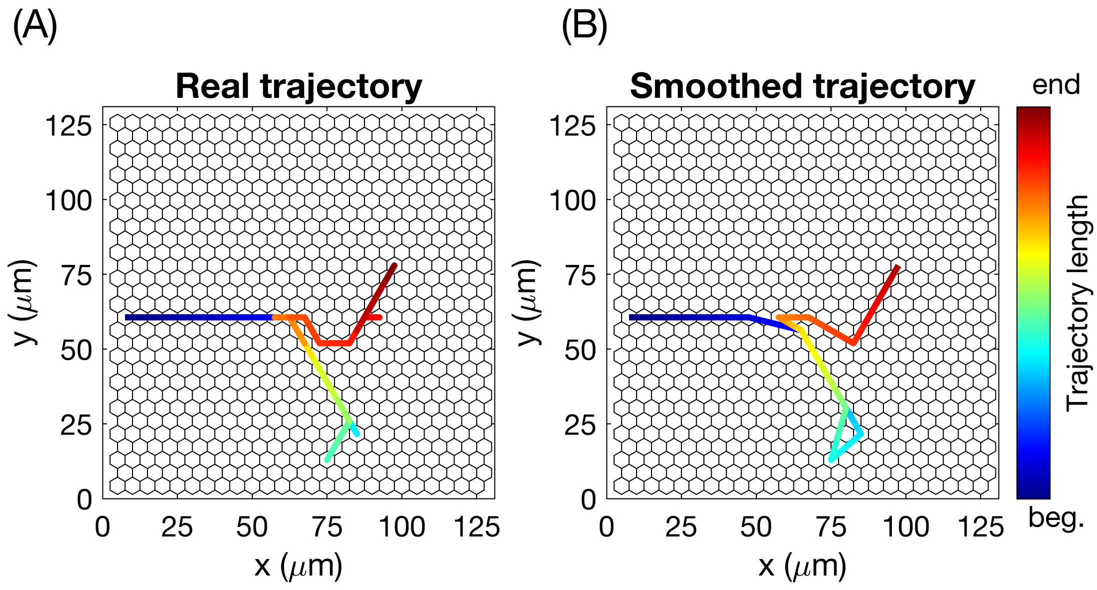


**SM-Fig 6. Illustrations of real and smoothed trajectories for an individual cell in a single realization. (A)** Unmodified (real) trajectory of an individual cell with a label, $\iota$, in a single realization, $p(\iota, t)$, $t \in [0, T_{max}]$. **(B)** The smoothed trajectory is obtained from the real trajectory, $p(\iota, t)$, by extracting a sample $p(\iota, t_k)$ such that $t_{k+1} - t_k \approx \Delta_{orient}$. The colourbar indicates the trajectory length.

**Algorithm 3.** Pseudocode algorithm for computing the directionality statistic.

Specify length of a time interval to define 'no movement' events, $\Delta_{still}$.
Introduce counters for anterograde movement, $count_a = 0$, retrograde movement, $count_r = 0$, and 'no movement' events, $count_s = 0$.
**for** each realization **do**
  Nullify the simulation time, $t = 0$.
  Create an empty array, $A = \varnothing$, to record labels of cells which moved during $\Delta_{still}$.
  $k = 1$.
  **while** $t < T_{max}$ **do**
    Obtain time step, $\tau$, for the next migration event, $\omega(i \to j)$, (see Algorithm 1).
    Identify labels of cells whose nuclei are in voxels $v_i$ and $v_j$, $\iota_i$ and $\iota_j$, respectively, (if $E_j(t) = 0$, i.e. $v_j$ is empty, then only $\iota_i$).
    $count_a = count_a + 1$.
    $A = A \bigcup \{\iota_i\}$.
    **if** $E_j(t) = 1$ **then**
      $count_r = count_r + 1$.
      $A = A \bigcup \{\iota_j\}$.
    **end if**
    Perform the migration event.
    $t = t + \tau$.
    **if** $t > \Delta_{still} k$ **then**
      $count_s = count_s + (max\_cell - \text{Size}(\text{Unique}(A)))$, where $max\_cell = \sum_{i \in \mathcal{I}} E_i(t)$.
      $A = \varnothing$.
      $k = k + 1$.
    **end if**
  **end while**
**end for**
The output is given by three counters for each type of cell movement: $count_a$, $count_r$ and $count_s$, which we use to make histograms of the directionality statistic (as in Fig 10(E) of the main text).

In Algorithm 3, routine Unique(·) returns an array of unique cell labels in a given set; routine Size(·) returns the cardinal of a given set.

**Tip cell proportion** This metric, $\mathcal{R}_{tips}(t)$, is defined as the ratio of cells with tip cell phenotype to the total number of cells in the system at a predetermined time, $t$. To compute its average over a number of realisations, we define a partitioning of the simulation time interval, $\mathcal{T}_{tips} = \mathcal{T}(\Delta_{tips})$, where $\Delta_{tips}$ is a uniform partitioning step.

We recall that we use the parameter characterising the baseline gene expression of Delta, $b_D$, as a threshold to define a cell's phenotype (see Eq (3) in the main text). Thus, $\mathcal{R}_{tips}(t)$ in a single realisation is computed as follows

$$\mathcal{R}_{tips}(t) = \frac{\text{Number of tip cells}}{\text{Total number of cells}} = \frac{\sum\limits_{i\in\mathcal{I}:\ D_i(t)\geqslant b_D} E_i(t)}{\sum\limits_{i\in\mathcal{I}} E_i(t)}. \tag{9}$$

The pseudocode for computing the tip cell proportion statistic, $\mathcal{R}_{tips}(t)$, is shown in Algorithm 4.

**Algorithm 4.** Pseudocode algorithm for computing the tip cell proportion statistic.

```
Specify the partitioning of the time interval, T_tips.
Nullify the realisation counter, r = 0.
while r < N_R do
    r = r + 1.
    Nullify the simulation time, t = 0.
    Set k = 0.
    while t < T_max do
        Obtain time step, τ, for the next migration event (see Algorithm 1).
        if t < t_k < t + τ then
            Compute R^r_tips(t_k) as in Eq (9).
            k = k + 1.
        end if
    end while
end while
Average the statistic over the performed realisations, R_tips(t_k) = (1/N_R) Σ_{r=0}^{N_R} R^r_tips(t_k),
for each k = 0, ..., K.
```

### 6 Mixing measure

A general definition of the mixing measure, $\mathcal{M}(t)$, is given by Eq (22)-(23) of the main text (see also Fig 6 of the main text for an illustration). In order to fully determine $\mathcal{M}(t)$, we need to specify the distance function, $d(\cdot,\cdot,\cdot)$. Since ECs migrate within empty sleeves of vascular guidance tunnels created due to ECM proteolysis [10], we convert the simulated network into a directed graph based on the configuration of the ECM-free tunnels (defines the set of graph vertices) and the ECM fibril orientation (defines the set of edges). Pairwise distances between cells are thus computed as the shortest possible paths within the graph. Specifically, we use the classical Dijkstra algorithm with Manhattan distance function [23]. Likewise, the maximum distance, $d_{max}$, used as a normalisation constant in Eq (22) of the main text, is the maximum Dijkstra distance in the generated graph.

We now proceed to provide a more detailed description of the numerical procedure used to convert a simulated vascular network into a graph, compute distances between the vertices of this graph, define a set of indices, $\mathcal{I}_{cluster}$, and compute the temporal evolution of the mixing measure in a single numerical realization.

#### Generating a graph from a simulated network

The algorithm for converting a simulated vascular network into a directed graph, $\mathcal{G}(\mathcal{N}, \mathcal{E})$, is based on the following.

I. After reaching the final simulation time, $T_{max}$, in a single realization, we consider the final state of the distribution of ECM concentration, $\boldsymbol{c}$. Based on this quantity, we determine the voxels that were explored by cell migration (thus generating the vascular guidance tunnels). Initially $c_i(t = 0) = c_{max}$ for $i \in \mathcal{I} \backslash \mathcal{I}_{init}$ (see S3 Table). Since during cell migration the ECM is degraded (see Eq (16) in the main text), voxels $v_i$, which have been visited by a cell are such that $c_i(t = T_{max}) < c_{max}$. This allows us to define the so-called explored network, $\mathcal{N}$, as a subset of voxel indices such that

$$\mathcal{N} = \{i \in \mathcal{I} \text{ such that } c_i < c_{max}\}\,. \tag{10}$$

The voxels with indices in $\mathcal{N}$ is the set of vertices in the graph, $\mathcal{G}(\mathcal{N}, \mathcal{E})$, to be constructed. Instead of using 2D indices, for simplicity, we label them in an arbitrary order.

II. To determine the set of edges, $\mathcal{E}$, we look at the configuration of the ECM fibril orientation, $\boldsymbol{l}$ (orientation landscape (OL) variable). Vertices corresponding to the voxels $v_i$ and $v_j$, $i, j \in \mathcal{N}$ are connected if the OL of voxel of origin, $l_i = \{l_i^{\bar{s}}\}_{\bar{s} \in \mathcal{S}}$, possesses a component $s \in \mathcal{S}$ greater than the initialisation value, $\Delta_{init}$. Here $s$ is a unit vector connecting the centres of $v_i$ and $v_j$ (see SM-Fig 7). The rationale for this rule is that this condition implies that the direction $s$ has been explored by a migration event from voxel $v_i$ to $v_j$ during simulation (see Eq (16) in the main text). Mathematically, it reads

   (i) there is an edge $e_{ij}$, $i, j \in \mathcal{N}$ if $l_i^s > \Delta_{init}$ $s = h^{-1}(q_j - q_i) \in \mathcal{S}$.

   In the example shown in SM-Fig 8, condition (i) is satisfied for the pairs of vertices with indices $1 \to 2$ and $3 \to 4$.

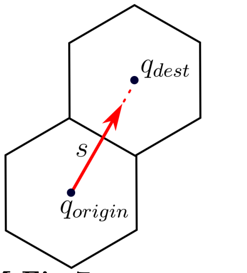


**SM-Fig 7. Illustration of the direction vector, $s$.** $s = h^{-1}(q_{dest} - q_{origin})$, where $q_k \in \mathbb{R}^2$ denotes a vector of coordinates of the centre of a voxel with index $k$ and $h$ is the voxel width.

III. In most simulated networks, we find sprouts of width of more than one cell (for example, as shown in SM-Fig 8(A)). To avoid infinite distances between first-neighbour voxels which belong to the same sprout and do not point towards each other but rather are aligned in the same direction, we connect these vertices if the corresponding voxels exhibit the same explored direction, $\bar{s}$. This reads as follows

   (ii) there is an edge $e_{ij}$, $i, j \in \mathcal{N}$ if there exists $\bar{s} \in \mathcal{S}$ such that $l_i^{\bar{s}} > \Delta_{init}$ and $l_j^{\bar{s}} > \Delta_{init}$.

   In SM-Fig 8(A), voxels (vertices) 1 and 3 are neighbouring, but do not point towards each other. Nonetheless, in a generated graph (SM-Fig 8(B)) they are connected, since they both have explored the rightward OL direction ($\bar{s} = r$). The same argument holds for the pairs of vertices $2 \leftrightarrow 4$, $1 \leftrightarrow 4$, $2 \to 1$ and $4 \to 3$.

IV. We assume that all edges of the graph have the same weight equal to unity.

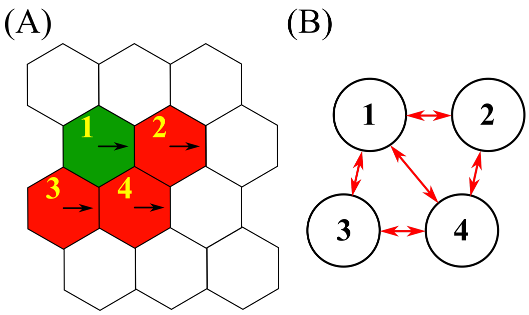


**SM-Fig 8.** **(A)** A simple example of a simulated network with $|\mathcal{N}| = 4$. The green colour corresponds to tip cell phenotype, red colour to stalk phenotype. Since there are cells in these voxels, the ECM concentration is less than $c_{max}$. Thus, these voxels form the explored network, $\mathcal{N}$ (numbering is done in an arbitrary fashion). Explored OL variable directions are indicated by arrows. **(B)** The graph corresponding to the simulated network from **(A)**.

We thus formulate a general algorithm, Algorithm 5, for converting a simulated vascular network into a graph.

**Algorithm 5.** Pseudocode algorithm to generate a graph from a simulated network.

**function** NETWORK TO GRAPH ( $\boldsymbol{c}$, $c_{max}$, $\boldsymbol{l}$, $\Delta_{init}$, $\mathcal{I}$, $\mathcal{S}$)
    Create set of graph vertices $\mathcal{N} \leftarrow \{i \in \mathcal{I}$ such that $c_i < c_{max}\}$ ▷ I.
    Create an empty set of edges $\mathcal{E} \leftarrow \{\ \}$
    **for** $i \in \mathcal{N}$ **do**
        **for** $s \in \mathcal{S}$ **do**
            $j \in \mathcal{I}$ such that $q_j = q_i + hs$ ▷ SM-Fig 7
            **if** such $j$ does not exist or $j \notin \mathcal{N}$ **then**
                **continue**
            **end if**
            **if** there exists $\bar{s} \in \mathcal{S}$ such that either $\left[l_i^{\bar{s}} > \Delta_{init}\ \&\ \bar{s} = s\right]$ or $\left[l_i^{\bar{s}} > \Delta_{init}\ \&\ l_j^{\bar{s}} > \Delta_{init}\right]$ **then**
                Add an edge from vertex $i$ to $j$, $e_{i,j}$, to the set of edges $\mathcal{E}$ ▷ II., III., SM-Fig 8
            **end if**
        **end for**
    **end for**
    Create a graph $\mathcal{G}(\mathcal{N}, \mathcal{E})$ with all edge costs equal to 1 ▷ IV.
    **return** $\mathcal{G}(\mathcal{N}, \mathcal{E})$
**end function**

We illustrate Algorithm 5 step by step with a simple example based on the small simulated vascular network in SM-Fig 9. The final configuration of the explored network is shown in SM-Fig 9(A) (explored vascular guidance tunnels with collagen concentration less than 1.0). The OL configuration is shown by arrows on this plot. The voxels corresponding to the explored network, $\mathcal{N}$, are then labelled by numbers in SM-Fig 9(B). They form the set of vertices of the graph. Applying Algorithm 5 to it, the simulated network is transformed into a graph (see SM-Fig 9(C)-(D), all edges are bidirectional).

#### Distances in a graph

The output of Algorithm 5 is an unweighted directed graph $\mathcal{G}(\mathcal{N}, \mathcal{E})$. We can use it to compute a matrix of shortest distances in a graph for each pair of vertices (i.e. explored voxels) in $\mathcal{N}$. To do so, we use the classical Dijkstra algorithm [23]. This algorithm takes as input a directional graph with non-negative edge weights (for example, the one shown in SM-Fig 9(D)) and computes a matrix of shortest paths between each pair of vertices, using edge costs. The distance of a path is calculated simply by adding up the weights of

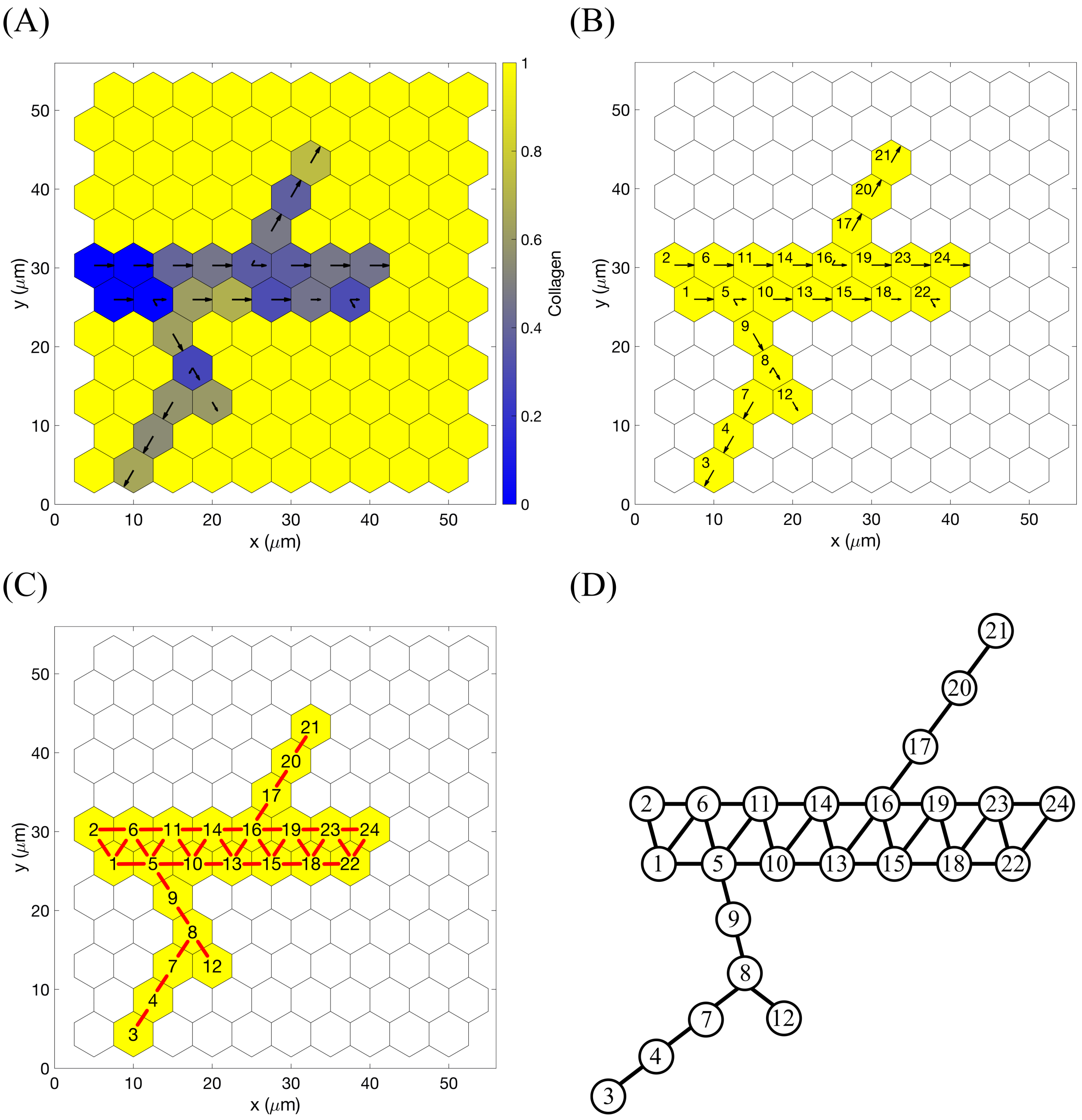


**SM-Fig 9. An example of converting a simulated network into a graph. (A)** The final configuration of the ECM concentration, $\boldsymbol{c}$. Here $c_{max} = 1.0$, thus the explored network, $\mathcal{N}$, corresponds to all voxels $v_i$ with $c_i < 1.0$. The colour bar indicates collagen concentration. Arrows correspond to the explored OL configuration (ECM fibril alignment). **(B)** Labelling (in an arbitrary fashion) the explored network, $\mathcal{N}$ (see Eq (10)). Voxels, indices of which belong to it, are coloured in yellow. **(C)** Applying Algorithm 5 to generate a directed graph from the network. Edges between vertices are shown in red (all bidirectional). **(D)** The final graph (all edges are bidirectional, arrows are omitted for simplicity) that is provided as an input to the Dijkstra algorithm to compute shortest paths between graph vertices. All edges have equal weights of 1.

all edges constituting the path. Continuing with the example from SM-Fig 9, the matrix of shortest path distances for the graph from SM-Fig 9(D) is shown in SM-Table 3. The quantity $d_{max}$ in Eq (23) in the main text is then the maximum value in this matrix.

| | | | | | | | | | | | | | | | | | | | | | | | |
|---|---|---|---|---|---|---|---|---|---|---|---|---|---|---|---|---|---|---|---|---|---|---|---|
| 0 | 1 | 6 | 5 | 1 | 1 | 4 | 3 | 2 | 2 | 2 | 4 | 3 | 3 | 4 | 4 | 5 | 5 | 5 | 6 | 7 | 6 | 6 | 7 |
| 1 | 0 | 7 | 6 | 2 | 1 | 5 | 4 | 3 | 3 | 2 | 5 | 4 | 3 | 5 | 4 | 5 | 6 | 5 | 6 | 7 | 7 | 6 | 7 |
| 6 | 7 | 0 | 1 | 5 | 6 | 2 | 3 | 4 | 6 | 6 | 5 | 7 | 7 | 8 | 8 | 9 | 9 | 9 | 10 | 11 | 10 | 10 | 11 |
| 5 | 6 | 1 | 0 | 4 | 5 | 1 | 2 | 3 | 5 | 5 | 3 | 6 | 6 | 7 | 7 | 8 | 8 | 8 | 9 | 10 | 9 | 9 | 10 |
| 1 | 2 | 5 | 4 | 0 | 1 | 3 | 2 | 1 | 1 | 1 | 3 | 2 | 2 | 3 | 3 | 4 | 4 | 4 | 5 | 6 | 5 | 5 | 6 |
| 1 | 1 | 6 | 5 | 1 | 0 | 4 | 3 | 2 | 2 | 1 | 4 | 3 | 2 | 4 | 3 | 4 | 5 | 4 | 5 | 6 | 6 | 5 | 6 |
| 4 | 5 | 2 | 1 | 3 | 4 | 0 | 1 | 2 | 4 | 4 | 2 | 5 | 5 | 6 | 6 | 7 | 7 | 7 | 8 | 9 | 8 | 8 | 9 |
| 3 | 4 | 3 | 2 | 2 | 3 | 1 | 0 | 1 | 3 | 3 | 1 | 4 | 4 | 5 | 5 | 6 | 6 | 6 | 7 | 8 | 7 | 7 | 8 |
| 2 | 3 | 4 | 3 | 1 | 2 | 2 | 1 | 0 | 2 | 2 | 2 | 3 | 3 | 4 | 4 | 5 | 5 | 5 | 6 | 7 | 6 | 6 | 7 |
| 2 | 3 | 6 | 5 | 1 | 2 | 4 | 3 | 2 | 0 | 1 | 4 | 1 | 1 | 2 | 2 | 3 | 3 | 3 | 4 | 5 | 4 | 4 | 5 |
| 2 | 2 | 6 | 5 | 1 | 1 | 4 | 3 | 2 | 1 | 0 | 4 | 2 | 1 | 3 | 2 | 3 | 4 | 3 | 4 | 5 | 5 | 4 | 5 |
| 4 | 5 | 5 | 3 | 3 | 4 | 2 | 1 | 2 | 4 | 4 | 0 | 5 | 5 | 6 | 6 | 7 | 7 | 7 | 8 | 9 | 8 | 8 | 9 |
| 3 | 4 | 7 | 6 | 2 | 3 | 5 | 4 | 3 | 1 | 2 | 5 | 0 | 1 | 1 | 1 | 2 | 2 | 2 | 3 | 4 | 3 | 3 | 4 |
| 3 | 3 | 7 | 6 | 2 | 2 | 5 | 4 | 3 | 1 | 1 | 5 | 1 | 0 | 2 | 1 | 2 | 3 | 2 | 3 | 4 | 4 | 3 | 4 |
| 4 | 5 | 8 | 7 | 3 | 4 | 6 | 5 | 4 | 2 | 3 | 6 | 1 | 2 | 0 | 1 | 2 | 1 | 1 | 3 | 4 | 2 | 2 | 3 |
| 4 | 4 | 8 | 7 | 3 | 3 | 6 | 5 | 4 | 2 | 2 | 6 | 1 | 1 | 1 | 0 | 1 | 2 | 1 | 2 | 3 | 3 | 2 | 3 |
| 5 | 5 | 9 | 8 | 4 | 4 | 7 | 6 | 5 | 3 | 3 | 7 | 2 | 2 | 2 | 1 | 0 | 3 | 1 | 1 | 2 | 4 | 3 | 4 |
| 5 | 6 | 9 | 8 | 4 | 5 | 7 | 6 | 5 | 3 | 4 | 7 | 2 | 3 | 1 | 2 | 3 | 0 | 1 | 4 | 5 | 1 | 1 | 2 |
| 5 | 5 | 9 | 8 | 4 | 4 | 7 | 6 | 5 | 3 | 3 | 7 | 2 | 2 | 1 | 1 | 1 | 1 | 0 | 3 | 4 | 2 | 1 | 2 |
| 6 | 6 | 10 | 9 | 5 | 5 | 8 | 7 | 6 | 4 | 4 | 8 | 3 | 3 | 3 | 2 | 1 | 4 | 3 | 0 | 1 | 5 | 4 | 5 |
| 7 | 7 | 11 | 10 | 6 | 6 | 9 | 8 | 7 | 5 | 5 | 9 | 4 | 4 | 4 | 3 | 2 | 5 | 4 | 1 | 0 | 6 | 5 | 6 |
| 6 | 7 | 10 | 9 | 5 | 6 | 8 | 7 | 6 | 4 | 5 | 8 | 3 | 4 | 2 | 3 | 4 | 1 | 2 | 5 | 6 | 0 | 1 | 1 |
| 6 | 6 | 10 | 9 | 5 | 5 | 8 | 7 | 6 | 4 | 4 | 8 | 3 | 3 | 2 | 2 | 3 | 1 | 1 | 4 | 5 | 1 | 0 | 1 |
| 7 | 7 | 11 | 10 | 6 | 6 | 9 | 8 | 7 | 5 | 5 | 9 | 4 | 4 | 3 | 3 | 4 | 2 | 2 | 5 | 6 | 1 | 1 | 0 |

**SM-Table 3.** The matrix of lengths of shortest paths between each pair of vertices in the graph shown in SM-Fig 9(D). Zero entries on the main diagonal indicate that there are no self-loops in the graph.

Using Algorithm 5 to generate a graph from the simulated network and the Dijkstra algorithm as a distance function between its vertices, we complete the definition of the mixing measure, $\mathcal{M}(t)$, Eq (22) of the main text.

#### Defining the time moments to compute the mixing statistic

To obtain the time evolution of the mixing measure averaged over a number of realisations, we need to partition the time interval of our simulations and compute $\mathcal{M}(\cdot)$ for each time instant belonging to the partition over all performed stochastic realisations. Two important issues need to be addressed:

I. $\mathcal{M}(t)$ is calculated as a normalised difference between the distance among cells in the cluster at some time, $t$, and the distance between the same cells at later time $(t + t_m)$. Thus, if $T_{max}$ is the final simulation time, then the last time moment for computing the mixing quantity is $(T_{max} - t_m)$.

II. Our simulations are stochastic. Consequently, time steps of events are not known a priori but rather sampled during each individual realisation. If we decide to partition the simulation time in some predetermined way for all realisations, $\mathcal{T} = \left\{\tau_k = \Delta_{mix} k,\ k = \overline{1, K},\ K = \left\lfloor \frac{T_{max} - t_m}{\Delta_{mix}} \right\rfloor \right\}$ (where $\Delta_{mix}$ is a uniform partitioning step), then we will not be able to calculate the mixing quantity at *exactly that* time instant. The time steps of the stochastic simulations are usually much smaller than the chosen partitioning time step $\Delta_{mix}$. Therefore, if the current simulation time is $t < \tau_k < T_{max} - t_m$ and next event will happen at time $\tau_k < \bar{t} < T_{max} - t_m$, then we compute the mixing measure at the time moment $\bar{t}$, i.e. as soon as the next partitioning time has been passed.

#### Defining the voxel cluster

The setup of the numerical simulations is such that we assume there is a set of voxel indices, $\mathcal{I}_{VP}$, corresponding to a vascular plexus from which the cells are migrating (see Appendix A). A Dirichlet boundary condition of a constant cell number is maintained at these voxels, i.e. if a cell migrates from one of these voxels, a new one immediately appears. Thus, choosing $\mathcal{I}_{VP} \subset \mathcal{I}_{cluster}$ guarantees that there are always cells present in these voxels, and the mixing measure makes sense at all times. However, $\mathcal{I}_{cluster}$ can be any set of indices of the lattice. Choosing a random position for the voxel cluster might lead to a situation when there are no cells present at the cluster location and we cannot compute the mixing quantity. We set $C = \mid \mathcal{I}_{cluster} \mid$, predetermined cluster size. In the main text, $\mathcal{I}_{cluster} = \mathcal{I}_{init}$ (**Setup 1** from S4 Table) was used, thus $C = 4$.

Whenever we decide to compute the mixing measure, we look at the labels of the cells located in the voxels in $\mathcal{I}_{cluster}$. Then we track position of these cells and compute the mixing quantity using Eqs (22)-(23) in the main text. $\mathcal{I}_{cluster}$ is fixed over all performed realisations, whereas labels of cells positioned in it for each desirable time moment change.

#### Mixing measure

In Algorithm 6 we lay out the general procedure for computing $\mathcal{M}(t)$ for a single realization. The mixing statistic is computed as a mean value of the mixing quantities for each moment of the time partition, $\mathcal{T}$, over all performed realisations.

**Algorithm 6.** Pseudocode algorithm for computing the mixing measure in a single stochastic realization of the model.

1: Define predetermined partitioning of the time interval $\mathcal{T} = \{\tau_k\}_1^K$.
2: Create two empty arrays of size $K \times C$ to record cell indices, $A_{id}$, (see SM-Fig 10(A)) and their final positions, $P$, (see SM-Fig 10(B)) for each time moment of the partition $\mathcal{T}$.
3: Start stochastic simulation of the multiscale model. Let $t$ denote the current simulation time.
4: Whenever simulation time $t > \tau_k$ for some $k$ that has not been computed yet, record in $A_{id}$ cell indices positioned at voxels with indexes $\mathcal{I}_{cluster}$, track positions of these cells and record their locations in the lattice at time $[t + t_m]$ and save them in the array $P$ (in the corresponding $k^{th}$ row).
5: When $t \geqslant T_{max}$ terminate the simulation.
6: Convert the final simulated network into a graph using Algorithm 5.
7: Compute the initial distance between cells in the cluster, $d_{init}$, that is the same for all $k$ (since cells are taken from the same lattice locations, $\mathcal{I}_{cluster}$), using the generated graph and vertices corresponding to $\mathcal{I}_{cluster}$ voxels.
8: For each $k$ in $\overline{1, K}$, compute the sum of the pair-wise distances in the graph between cells with indices from the $k^{th}$ row of $A_{id}$ taking their recorded final locations from the $k^{th}$ row of the array $P$. Denote this quantity as $d_k$.
9: Find the distance of the longest path in the graph, $d_{max}$, as the maximum entry in the matrix of Dijkstra shortest paths.
10: Compute the mixing quantity for each $k$ in $\overline{1, K}$ using Eq (22) of the main text which, in the notation of this algorithm, corresponds to $m_k = \frac{d_k - d_{init}}{C\ d_{max}}$.
11: Record in the output file the mixing quantities, $m_k$, for all $k = \overline{1, K}$.

**(A)**

$C = \mid \mathcal{I}_{cluster} \mid = \mid \{i_1, i_2, \ldots, i_C\} \mid$

$K$

| | | | |
|---|---|---|---|
| The index of a cell located at the voxel $v_{i_1}$, $i_1 \in \mathcal{I}_{cluster}$, at time $\tau_1$. | The index of a cell located at the voxel $v_{i_2}$, $i_2 \in \mathcal{I}_{cluster}$, at time $\tau_1$. | $\ldots$ | The index of a cell located at the voxel $v_{i_C}$, $i_C \in \mathcal{I}_{cluster}$, at time $\tau_1$. |
| The index of a cell located at the voxel $v_{i_1}$, $i_1 \in \mathcal{I}_{cluster}$, at time $\tau_2$. | The index of a cell located at the voxel $v_{i_2}$, $i_2 \in \mathcal{I}_{cluster}$, at time $\tau_2$. | $\ldots$ | The index of a cell located at the voxel $v_{i_C}$, $i_C \in \mathcal{I}_{cluster}$, at time $\tau_2$. |
| $\ldots$ | $\ldots$ | $\ldots$ | $\ldots$ |
| The index of a cell located at the voxel $v_{i_1}$, $i_1 \in \mathcal{I}_{cluster}$, at time $\tau_K$. | The index of a cell located at the voxel $v_{i_2}$, $i_2 \in \mathcal{I}_{cluster}$, at time $\tau_K$. | $\ldots$ | The index of a cell located at the voxel $v_{i_C}$, $i_C \in \mathcal{I}_{cluster}$, at time $\tau_K$. |

**(B)**

$C = \mid \mathcal{I}_{cluster} \mid = \mid \{i_1, i_2, \ldots, i_C\} \mid$

$K$

| | | | |
|---|---|---|---|
| Position of the cell with index $A_{id}[1, 1]$ at time $\tau_1 + t_m$. | Position of the cell with index $A_{id}[1, 2]$ at time $\tau_1 + t_m$. | $\ldots$ | Position of the cell with index $A_{id}[1, C]$ at time $\tau_1 + t_m$. |
| Position of the cell with index $A_{id}[2, 1]$ at time $\tau_2 + t_m$. | Position of the cell with index $A_{id}[2, 2]$ at time $\tau_2 + t_m$. | $\ldots$ | Position of the cell with index $A_{id}[2, C]$ at time $\tau_2 + t_m$. |
| $\ldots$ | $\ldots$ | $\ldots$ | $\ldots$ |
| Position of the cell with index $A_{id}[K, 1]$ at time $\tau_K + t_m$. | Position of the cell with index $A_{id}[K, 2]$ at time $\tau_K + t_m$. | $\ldots$ | Position of the cell with index $A_{id}[K, C]$ at time $\tau_K + t_m$. |

**SM-Fig 10. Array structures used in Algorithm 6. (A)** A schematic of the structure of the array of cell indices, $A_{id}$. **(B)** A schematic of the structure of the array of final cell positions, $P$.

### 7 Quantification of simulated vascular networks

To quantify in a rigorous way the branching structure of simulated vascular networks, we developed an algorithm to extract:

- vascular network area;

- number of branching points per 100 $\mu m^2$ area of vascular network;
- number of vessel segments;
- length of vessel segments.

We define a *vessel segment* as either a part of vascular network between two branching points or free sprouts, i.e. between a branching point and the leading edge of this sprout.

The orientation landscape variable and graph representation of the simulated network used to compute the mixing measure give us a direct way to perform vascular network quantification. Briefly, starting with the graph representation given by Algorithm 5 (see SM-Fig 11(B)), we reduce sprouts of width of more than one voxel to single-voxel sprouts. This gives us the 'skeleton' of the vascular network (see SM-Fig 11(C)). From this representation, it is straightforward to identify *branching points* as all vertices of degree greater than 2 and *vessel segments* are obtained by splitting the skeleton graph at the vertices of branching points (see SM-Fig 11(D)). We lay out the general procedure for extracting these statistics in Algorithm 7.

**Algorithm 7.** Pseudocode algorithm for extracting network quantification statistics in a single stochastic realization of the model.

Get graph representation of the vascular network, $\mathcal{G}(\mathcal{N}, \mathcal{E})$, from Algorithm 5.
Define a graph of the network skeleton, $\mathcal{G}_s(\mathcal{N}_s, \mathcal{E}_s) = \mathcal{G}(\mathcal{N}, \mathcal{E})$.
*Network area*, $A$, is equal to the number of vertices in the graph $\mathcal{G}_s$, $\mid \mathcal{N}_s \mid$, multiplied by the area of a single voxel ($0.5\sqrt{3}h^2$, in case of hexagonal voxel of width $h$ [$\mu m$]).
Create an empty set of pairs of vertices of $\mathcal{G}_s$ to be merged, $\mathcal{N}_{merge}$.
**for** $i \in \mathcal{N}$ **do**
    **for** $s \in \mathcal{S}$ **do**
        $j \in \mathcal{I}$ such that $q_j = q_i + hs$ ▷ SM-Fig 7
        **if** such $j$ does not exist or $j \notin \mathcal{N}$ **then**
            **continue**
        **end if**
        **if** there exists $\bar{s} \in \mathcal{S}$ such that $\left[l_i^{\bar{s}} > \Delta_{init} \text{ \& } l_j^{\bar{s}} > \Delta_{init}\right]$ **then**
            Add the pair of vertices $(i, j)$ to the set $\mathcal{N}_{merge}$
        **end if**
    **end for**
**end for**
**for** $(i, j) \in \mathcal{N}_{merge}$ **do**
    Merge vertex $i$ with vertex $j$, i.e. delete from $\mathcal{E}_s$ the edges $e_{i,j}$ and $e_{j,i}$ and all edges of type $e_{j,\cdot}$ ($e_{\cdot,j}$) become $e_{i,\cdot}$ ($e_{\cdot,i}$).
**end for**
Delete all vertices of degree 0 from $\mathcal{N}_s$.
Delete repeated edges in $\mathcal{G}_s(\mathcal{N}_s, \mathcal{E}_s)$.
Identify *branching points* as the set of vertices of degree greater than 2,
    $\mathcal{N}_{branch} = \{i \in \mathcal{N}_s \ s.t. \ deg(i) > 2\}$. ▷ SM-Fig 11(C)
*Number of branching points per 100 $\mu m^2$ of vascular network area*, $N_b = \frac{|\mathcal{N}_{branch}| \cdot 100 \mu m^2}{A}$.
The graph of vessel segments, $\mathcal{G}_{vessels}$, is obtained from $\mathcal{G}_s$ by duplicating each vertex in $\mathcal{N}_{branch}$ by its degree and splitting the graph, $\mathcal{G}_s$, at these points. ▷ SM-Fig 11(D)
*Number of vessel segments* is given by the number of connected components in the graph $\mathcal{G}_{vessels}$.
*Vessel segment lengths* are given by the number of edges in each connected component of $\mathcal{G}_{vessels}$ multiplied by the voxel width, $h$ [$\mu m$].

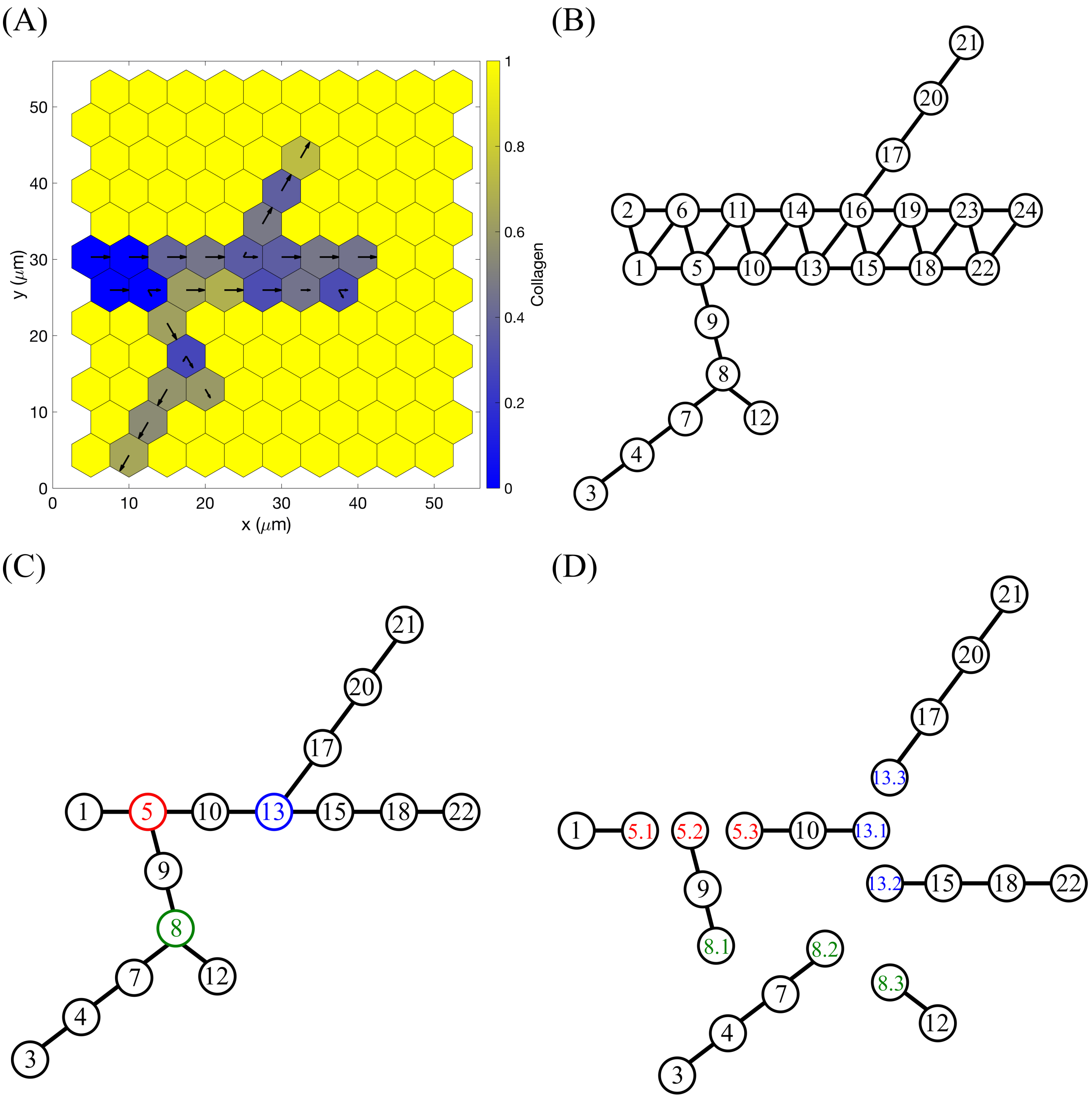


**SM-Fig 11. An example of branching structure quantification on a single simulated vascular network. (A)** The final configuration of the ECM concentration, $\boldsymbol{c}$, of a simulated vascular network. Arrows correspond to the explored OL configuration (ECM fibril alignment). **(B)** The graph representation of the vascular network from **(A)** obtained by applying to it Algorithm 5. **(C)** Vascular network 'skeleton', $\mathcal{G}_s$, obtained by reducing multiple-voxel vessels to single-voxel (see Algorithm 7). Branching points (defined as vertices of degree greater than 2) are highlighted by a different colour. **(D)** Splitting the network 'skeleton' at the branching points to obtain vessel segments graph, $\mathcal{G}_{vessels}$ (see Algorithm 7).

### 8 Sensitivity analysis

Since our subcellular scale model was based on the previous works [13, 14, 15, 16], we consider the parameters of this scale to be fixed (see S1 Table). By contrast, our implementation of cell migration and cell-ECM interactions is novel. Thus, we performed an extensive sensitivity analysis for the parameters corresponding to the cellular and tissue scales of our model (listed in S2 Table).

We define the set of baseline parameter values as

$$\bar{p} = \{K, k_m, k_D, \eta_{max}, s_c, D_c, \gamma_{max}, s_m, D_m, p_{max}, s_p, D_p, E_{F1}, E_{F2}, s_{F1}, s_{F2}, R_c, D_\omega, \eta_l, c_{max}, a, \Delta_l\}.$$

Brief interpretations of these parameters and references to the corresponding equations are given in SM-Table 4. Baseline parameter values, calibrated by comparing our model to experimental data from [1, 2, 24], are listed in S2 Table.

| Parameters | Interpretation | Reference |
|---|---|---|
| $K$, $k_m$, $k_D$ | cell exploratoriness | Eq (14) |
| $\eta_{max}$, $s_c$, $D_c$ | ECM proteolysis | Eqs (18)-(19) |
| $\gamma_{max}$, $s_m$, $D_m$ | BM assembly | Eqs (20)-(21) |
| $p_{max}$, $s_p$, $D_p$ | probability of cell overtaking | Eq (10)-(11) |
| $E_{F1}$, $E_{F2}$, $s_{F1}$, $s_{F1}$, $s_{F2}$ | cell-cell interaction | Eq (9) |
| $R_c$ | cellular scale cell-cell interaction radius | Table 1, $\boldsymbol{E^N}$ |
| $D_\omega$ | diffusion coefficient | Eq (7) |
| $\eta_l$ | ECM relaxation rate | Eq (17) |
| $c_{max}$ | maximum ECM concentration | Eq (8) |
| $a$ | polarity vector parameter | Eq (13) |
| $\Delta_l$ | orientation landscape update increment | Eq (16) |

**SM-Table 4. Parameters of the cellular and tissue scales used for sensitivity analysis** Equation references are for the main text.

We define $p_i$ as a vector of parameter values as in $\bar{p}$ except for the parameter at position $i$ which is incremented (or decremented) as indicated. For each of these parameter sets, we perform 100 realisations of our model using simulation **Setup 1** from S4 Table. The parameters of the subcellular scale are fixed for all experiments (see S1 Table). From these data we extract the following quantification metrics:

- anterograde cell proportion (directionality statistic);
- orientation;
- displacements;
- number of branching points per 100 $\mu m^2$ of vascular network area;
- number of vessel segments;
- vessel segment lengths.

In order to compare these metrics for modified parameter sets, $p_i$, to the baseline parameters, $\bar{p}$, we compute:

- Kolmogorov-Smirnov statistic:
$$\mathcal{D}^* = \max_x \mid \hat{F}_{\bar{p}}(x) - \hat{F}_{p_i}(x) \mid$$
Here $\hat{F}_{\bar{p}}(x)$ and $\hat{F}_{p_i}(x)$ are empirical cumulative distribution functions corresponding to the chosen metric for the sets of parameters $\bar{p}$ and $p_i$, respectively. $\mathcal{D}^* \in [0, 1]$, where values close to 0 indicate that the corresponding cumulative distributions are similar and values close to 1 indicate that $\hat{F}_{\bar{p}}(x)$ and $\hat{F}_{p_i}(x)$ differ significantly.

- Relative mean change:
$$\frac{\mu_{p_i} - \mu_{\bar{p}}}{\mu_{\bar{p}}},$$
where $\mu_{p_i}$ and $\mu_{\bar{p}}$ are the mean values of the chosen metric for $\bar{p}$ and $p_i$, respectively.

- Relative change of standard deviation:
$$\frac{\sigma_{p_i} - \sigma_{\bar{p}}}{\sigma_{\bar{p}}},$$
where $\sigma_{p_i}$ and $\sigma_{\bar{p}}$ are the standard deviations of the chosen metric for $\bar{p}$ and $p_i$, respectively.

We performed experiments for $\pm 0.1\%$, $\pm 1\%$, $\pm 5\%$, $\pm 5\%$, $\pm 10\%$, $\pm 15\%$ and $\pm 20\%$. The results for $\pm 10\%$ change in the parameter values are shown in SM-Figs 12, 13, 14 for the Kolmogorov-Smirnov statistic, relative change of means and relative change of standard deviation, respectively. From these plots, it can be seen that cell behaviour (such as anterograde cell proportion, orientation of cell trajectory and cell displacements) and branching structure of the simulated networks (number of branching points, number and length of vessel segments) are affected the most by variations in parameters $D_c$ and $D_m$ characterising ECM proteolysis and BM deposition rates, respectively. Less significant change in the metrics is seen for cell exploratoriness parameters, $K$ and $k_D$, and cell-cell adhesion parameters, $E_{F1}$ and $E_{F2}$. We also note that individual cell behaviour is less affected than the overall vascular network structure.

The results of the sensitivity analysis for $\pm 10\%$ for the final value of mixing measure (the last value in the time evolution of the mixing measure extracted from the numerical simulations, $\mathcal{M}(t - t_m)$) are shown in SM-Fig 15. This metric is more robust than the metrics considered in SM-Figs 12, 13, 14. Interestingly, variations in majority of parameters (both positive and negative) induce a decrease in the final value of the mixing measure.

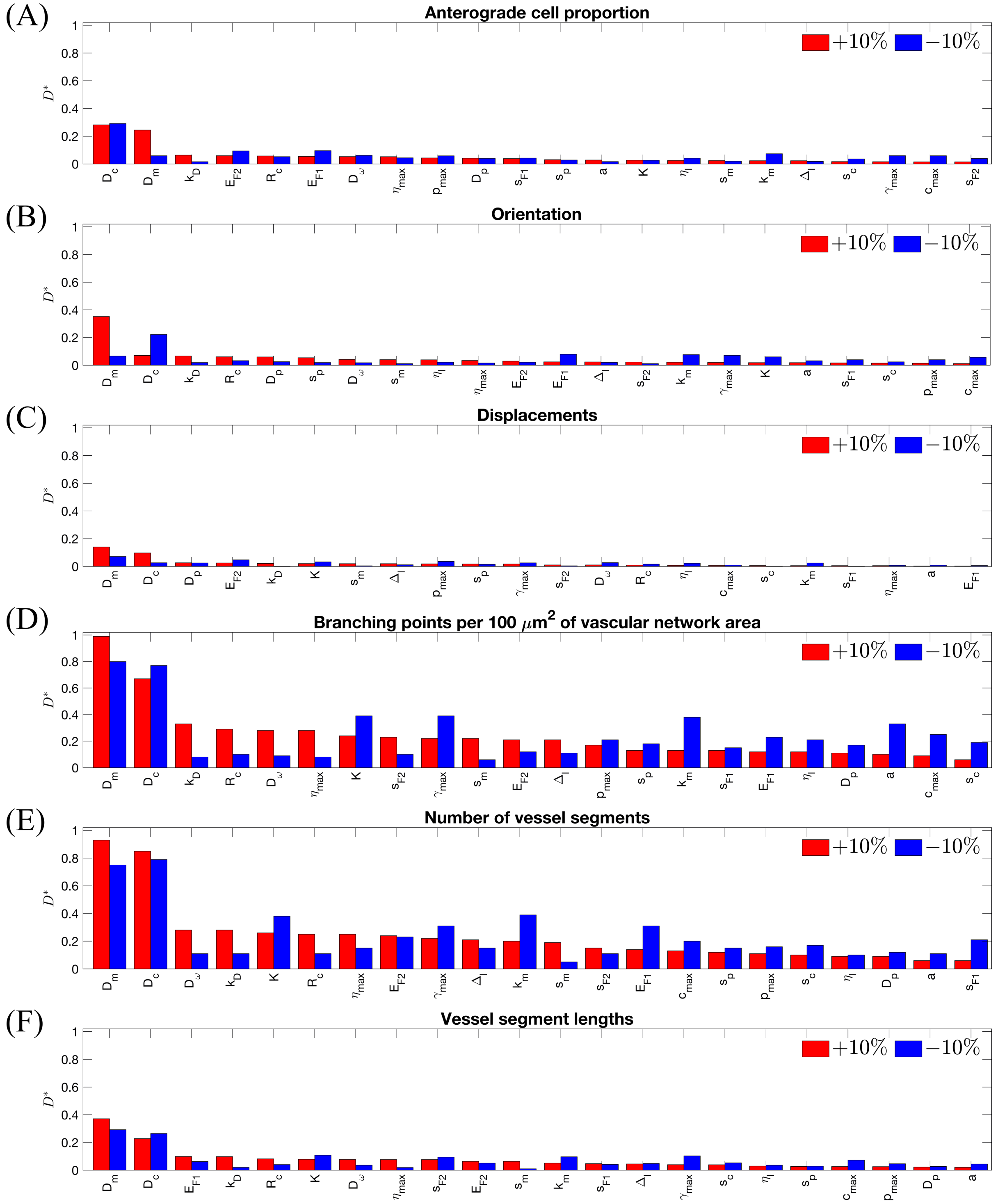


**SM-Fig 12. Sensitivity analysis results for Kolmogorov-Smirnov distance for the following metrics: (A)** anterograde cell proportion, **(B)** orientation, **(C)** displacements, **(D)** number of branching points per 100 $\mu m^2$ of vascular network area, **(E)** number of vessel segments, **(F)** vessel segment lengths. Simulations were performed using **Setup 1** from S4 Table and the parameter values as listed in S2 Table except for the parameter indicated at x-axis which was incremented (decremented) by 10%. For each set of parameters 100 realisations were performed.

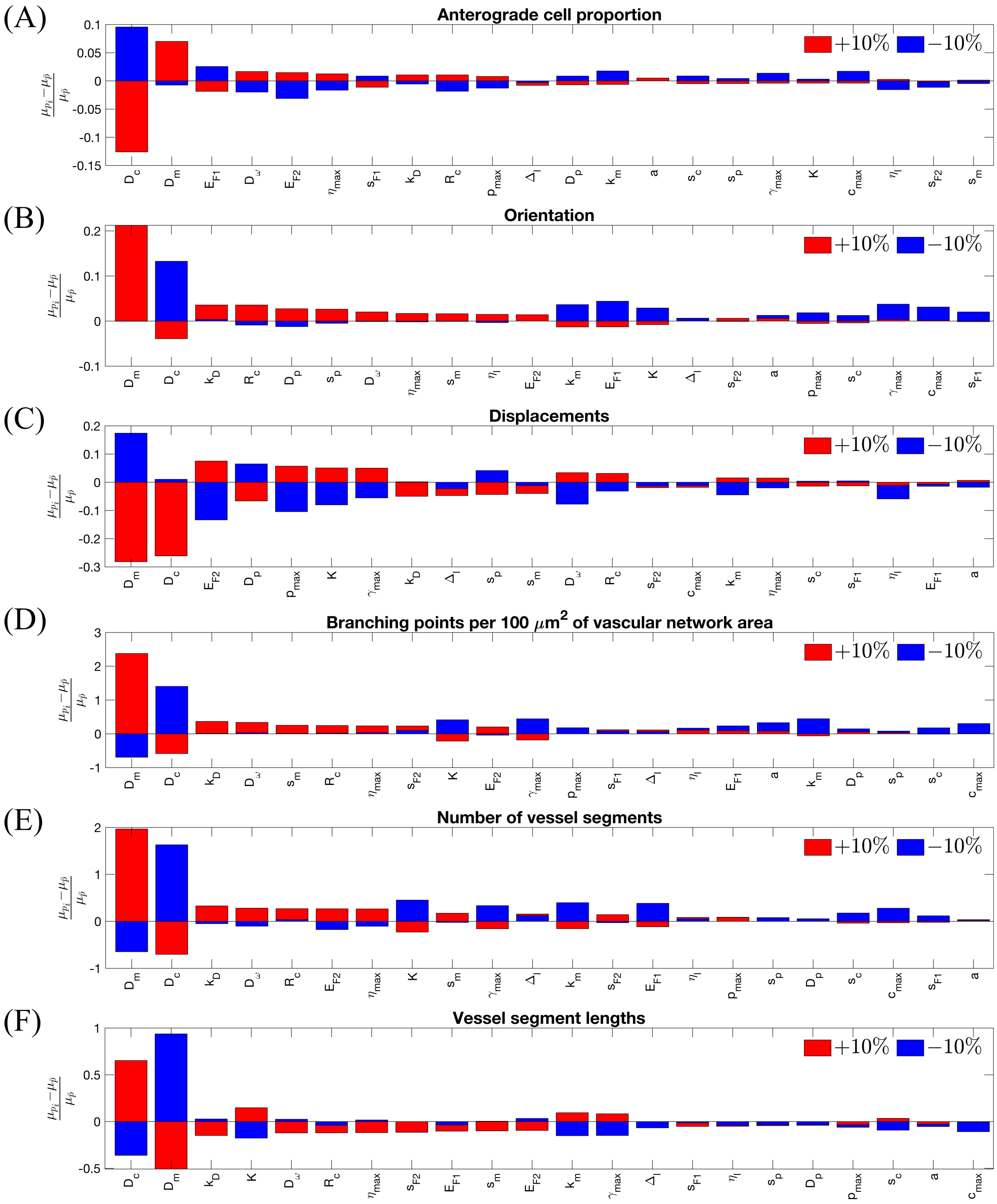


**SM-Fig 13. Sensitivity analysis results for relative mean change for the following metrics:** **(A)** anterograde cell proportion, **(B)** orientation, **(C)** displacements, **(D)** number of branching points per 100 $\mu m^2$ of vascular network area, **(E)** number of vessel segments, **(F)** vessel segment lengths. Simulations were performed using **Setup 1** from S4 Table and the parameter values as listed in S2 Table except for the parameter indicated at x-axis which was incremented (decremented) by 10%. For each set of parameters 100 realisations were performed.

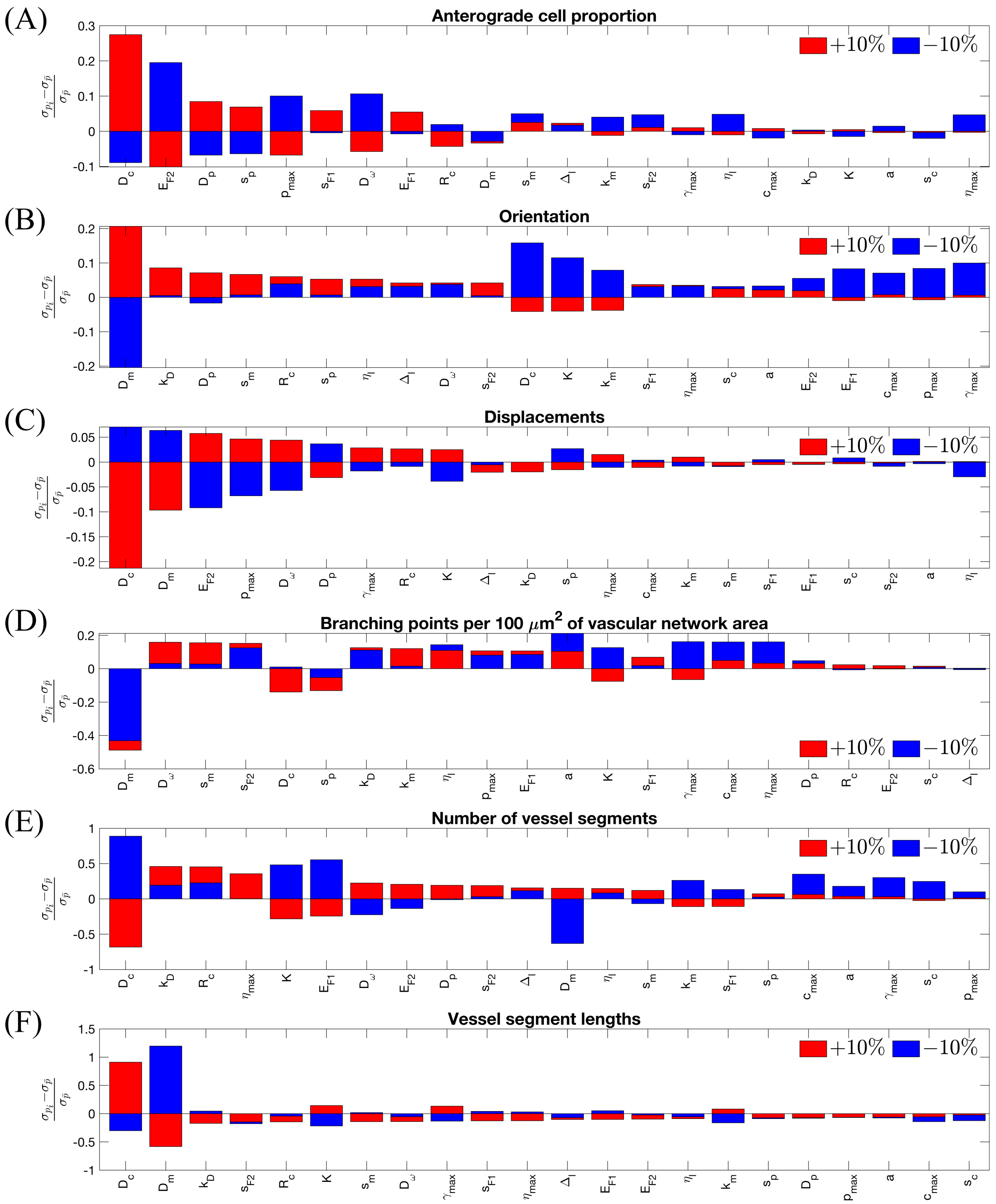


**SM-Fig 14. Sensitivity analysis results for relative change of standard deviation for the following metrics: (A)** anterograde cell proportion, **(B)** orientation, **(C)** displacements, **(D)** number of branching points per 100 $\mu m^2$ of vascular network area, **(E)** number of vessel segments, **(F)** vessel segment lengths. Simulations were performed using **Setup 1** from S4 Table and the parameter values as listed in S2 Table except for the parameter indicated at x-axis which was incremented (decremented) by 10%. For each set of parameters 100 realisations were performed.

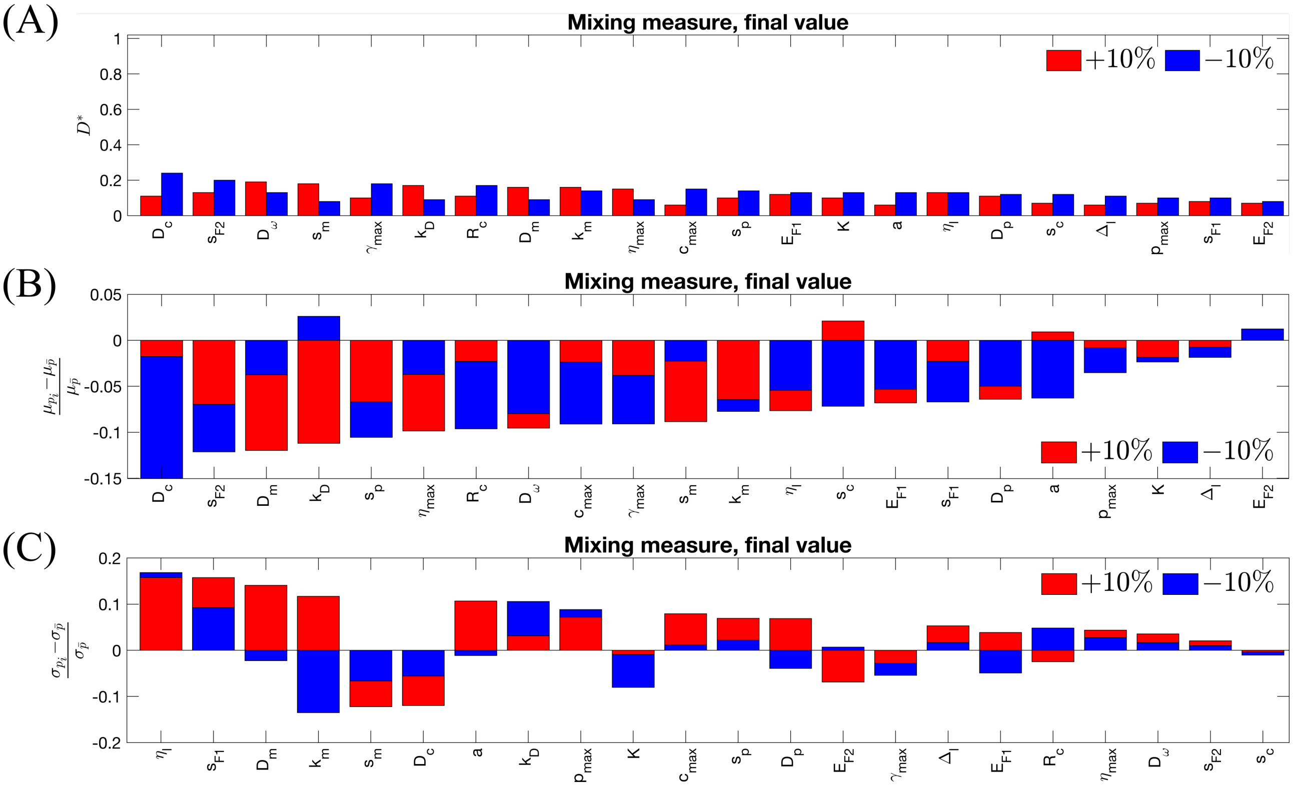


**SM-Fig 15. Sensitivity analysis results for final value of mixing measure: (A)** Kolmogorov-Smirnov distance, **(B)** relative mean change, **(C)** relative change of standard deviation. Simulations were performed using **Setup 1** from S4 Table and the parameter values as listed in S2 Table except for the parameter indicated at x-axis which was incremented (decremented) by 10%. For each set of parameters 100 realisations were performed.
