## Supplementary material for "A multiscale model of complex endothelial cell dynamics in early angiogenesis": S1 Appendix

### S1 Appendix. Computational simulations

**Model geometry** All simulations were performed on a rectangular lattice, $\mathcal{L} = \{v_i,\ i = (i_x, i_y)^T,\ i_x = 1, ..., N_I^x,\ i_y = 1, ..., N_I^y\}$, where $v_i$ stands for voxel indexed by $i$, and $i$ denotes the position of the voxel $v_i$ within the lattice, $\mathcal{L}$. The total voxel number $N_I = N_I^x N_I^y$. $N_I^x$ and $N_I^y$ vary for each type of numerical experiment described in S4 Table. The non-dimensional voxel width, $h = 0.04$, corresponds to 5 $\mu m$ (see S1 Text for details).

**Boundary conditions** Let $\mathcal{I}_B$ denote the set of voxels of $\mathcal{L}$ situated on its boundary, i.e.

$$\mathcal{I}_B = \{(1, i_y)^T,\ i_y = 1, ..., N_I^y\} \cup \{(N_I^x, i_y)^T,\ i_y = 1, ..., N_I^y\} \cup \{(i_x, 1)^T,\ i_x = 1, ..., N_I^x\} \cup \{(i_x, N_I^y)^T,\ i_x = 1, ..., N_I^x\}.$$

As mentioned before, we assume that our simulations take place on a time scale such that cell proliferation is negligible and sprout elongation is driven by cell migration from the initial vascular plexus (imitating an underlying vascular bed in *in vivo* or a cell implant in *in vitro* experiments). This is implemented as a Dirichlet boundary condition for the nucleus distribution variable, $\boldsymbol{E}$, for the set of voxels corresponding to the position of the vascular plexus, $\mathcal{I}_{VP}$,

$$E_i = 1\ \forall i \in \mathcal{I}_{VP},\ \forall t \geq 0.$$

The set $\mathcal{I}_{VP}$ for each numerical experiment is listed in S4 Table. When a cell migrates from a voxel belonging to $\mathcal{I}_{VP}$, a new cell is put in this voxel with the baseline expression of the subcellular scale variables (its phenotype is established according to its environment in the following simulation of the subcellular model).

For the voxels on the boundary, a no-flux boundary condition is assumed for cells: migration transitions to leave the domain are set to zero, $\omega(i \to \text{ exit } \mathcal{L}) = 0$, for $i \in \mathcal{I}_B$.

Since we assume that cells cannot leave the domain, we set the orientation landscape variable components pointing outside $\mathcal{L}$ to zero. Mathematically, let $n_e$ denote an external normal to $\mathcal{L}$, then

$$l_i^s = 0, \ \forall i \in \mathcal{I}_B \text{ and } \forall s \in \mathcal{S} \text{ s.t. } (s, n_e) = 1.$$

Here $(\cdot, \cdot)$ is the scalar product.

The rest of the variables, namely, the variables of the subcellular scale, ECM and BM components, $\boldsymbol{c}$ and $\boldsymbol{m}$, respectively, do not require any specific boundary conditions.

**Initial conditions** Let $\mathcal{I}_{init}$ denote the voxel indices of the initial cell positions, i.e. $E_i = 1$ for $i \in \mathcal{I}_{init}$ and $E_i = 0$, otherwise, at time $t = 0$. Given this set of indices, the variables are initialised as shown in S3 Table.

**Setups** We performed several types of simulation experiments. For each type we specify the lattice dimensions, $N_I^x$ and $N_I^y$, the set of indices corresponding to the vascular plexus, $\mathcal{I}_{VP}$, the initial cell nucleus positions, $\mathcal{I}_{init}$, the initial polarisation direction, $s_{init}$, the initial ECM and BM concentrations, $c_{init}$ and $m_{init}$, respectively, and the distribution of VEGF, $V$. The details are given in S4 Table.

**Simulations with mutant cells** Some of the simulation experiments were performed with mutant cells with modified gene expression of VEGFR1 and VEGFR2, with the aim of imitating the behaviour of those used in experiments reported in [7]. To do so, we modify some of the parameters of the subcellular VEGF-Delta-Notch signalling. The details are given below in Table A.

| **Name [7]** | **Shortened name** | **Description** | **Change in parameters** |
|---|---|---|---|
| VEGFR2$^{+/egfp}$ | VEGFR2$^{+/-}$ | Mutant cells heterozygous for VEGF Receptor 2 having half of the amount of VEGFR2 compared with the WT cells. | $b_{R2}^+ = 0.5b_{R2}$ |
| VEGFR2$^{+/egfp}$-DAPT | VEGFR2$^{+/-}$-DAPT | VEGFR2$^{+/-}$ mutant cells additionally exposed to DAPT, a $\gamma$-secretase inhibitor abolishing the Notch signalling. | $b_{R2}^+ = 0.5b_{R2}$, $I = 0$ |
| VEGFR1$^{+/lacz}$ | VEGFR1$^{+/-}$ | Mutant cells heterozygous for VEGF Receptor 1 having half of the amount of VEGFR1 compared with the WT cells. | $k_v^+ = 2k_v$ |
| VEGFR1$^{+/lacz}$-DAPT | VEGFR1$^{+/-}$-DAPT | VEGFR1$^{+/-}$ mutant cells additionally exposed to DAPT, a $\gamma$-secretase inhibitor abolishing the Notch signalling. | $k_v^+ = 2k_v$, $I = 0$ |
| WT-DAPT | WT-DAPT | Wild-type (WT) cells treated with DAPT, a $\gamma$-secretase inhibitor abolishing the Notch signalling. | $I = 0$ |

**Table A. Description of mutant cells treated/untreated with DAPT ($\gamma$-secretase inhibitor) used in simulations.** Changed parameters for mutant cells have + in their superscript position, compared to the wild-type (WT) cell parameters with no such superscripts.

In particular, VEGFR2$^{+/-}$ mutant cells have down-regulated (by $\approx$50%, single allele mutants) gene expression of VEGFR2, thus we set its baseline expression to half of that for the WT (see Table A). VEGFR1$^{+/-}$ mutant cells are characterised by halved gene expression of VEGFR1. VEGFR1 is known to be a sink receptor for VEGF: it has higher affinity for binding VEGF but low kinase activity. Hence, it competes with VEGFR2 for binding to VEGF but it has a minor role in signal transmission. Because of this, we have not considered it in our subcellular model of phenotype selection, and we need to account for the VEGFR1$^{+/-}$ mutant in an effective way. Specifically, we assume that down-regulation of VEGFR1 can be accounted for by a higher affinity of VEGFR2 for binding to VEGF. We set $k_v^+ = 2k_v$ as shown in Table A. Furthermore, in some of the experiments by Jakobsson et al. [7], cells were treated with a potent $\gamma$-secretase inhibitor, DAPT, which completely abolishes Notch signalling (see Fig 3i in [7]). $\gamma$-secretase is a protease that carries out the second cleavage releasing the active NICD. Therefore, when exposed to DAPT, the Notch receptor and its ligand should have the same dynamics as without DAPT, only NICD is not being produced. To introduce DAPT into our simulations, we set $I = 0$ for all cells in the simulation (both WT and mutant), leaving all other parameters unchanged (see Table A).
