## Supplementary material for "A multiscale model of complex endothelial cell dynamics in early angiogenesis": S1 Table

| Parameter | Units | Description | Value used in simulations | Ref. |
|---|---|---|---|---|
| $R_s$ | $\mu m$ | Interaction radius. | 15 | estim., [9, 39] |
| $b_N$ | $molec \cdot time^{-1}$ | Baseline Notch receptor expression. | 500 | [72, 73] |
| $b_D$ | $molec \cdot time^{-1}$ | Baseline Delta ligand expression. | 800 | [72, 73] |
| $b_{R2}$ | $molec \cdot time^{-1}$ | Baseline VEGFR2 expression. | 800 | [73] |
| $I_0$ | $molec$ | Activation threshold for NICD. | 100 | [73] |
| $R2_0^*$ | $molec$ | Activation threshold for activated VEGFR2. | 200 | [73] |
| $\lambda_{I,N}$ | $dimensionless$ | Weight factor characterising fold change of the production rate of Notch receptor depending on the NICD concentration. | 4.0 | [72, 73] |
| $\lambda_{aR2,D}$ | $dimensionless$ | Weight factor characterising fold change of the production rate of Delta ligand depending on the activated VEGFR2 concentration. | 2.0 | [73] |
| $\lambda_{I,R2}$ | $dimensionless$ | Weight factor characterising fold change of the production rate of VEGFR2 depending on the NICD concentration. | 0.0 | [73] |
| $n_N$ | $dimensionless$ | Cooperativity parameter for Hill function for NICD-dependent Notch up-regulation. | 2 | [71] |
| $n_D$ | $dimensionless$ | Cooperativity parameter for Hill function for activated VEGF-dependent Delta up-regulation. | 1 | [71] |
| $n_{R2}$ | $dimensionless$ | Cooperativity parameter for Hill function for NICD-dependent VEGFR2 repression. | 1 | [71] |
| $V$ | $molec$ | External VEGF. | 2500 (Fig 3(E)); $0 - 2500$ (Fig 3(F)); $\{0, 2500, 25000\}$ (in the rest of the simulations) | [72, 73] |
| $D_{ext}$ | $molec$ | External Delta ligand. | $0 - 3000$ (Fig 3(E)-(F)); calculated from adjacent cells (in the rest of the simulations) | [72, 73] |
| $N_{ext}$ | $molec$ | External Notch receptor. | 1000 (Fig 3(E)-(F)); calculated from adjacent cells (in the rest of the simulations) | [72, 73] |
| $k_t$ | $molec^{-1} \cdot time^{-1}$ | Trans-binding rate for Notch receptor and Delta ligand. | $5.0e-5$ | [71] |
| $k_c$ | $molec^{-1} \cdot time^{-1}$ | Cis-interaction rate for Notch receptor and Delta ligand. | $6.0e-4$ | [71] |
| $k_v$ | $molec^{-1} \cdot time^{-1}$ | Binding rate for VEGFR2 and external VEGF. | $5.0e-5$ | [73] |
| $\eta$ | $dimensionless$ | Endocytic regulation of Notch signalling. | 0.5 | estim., [96] |
| $\gamma$ | $time^{-1}$ | Degradation rate of proteins. | 0.1 | [73] |
| $\gamma_e$ | $time^{-1}$ | Degradation rate of activated receptors. | 0.5 | [73] |

**S1 Table. Baseline parameter values for the VEGF-Delta-Notch subcellular model.** Description and reference values used in simulations of the subcellular VEGF-Delta-Notch signalling.
