## Supplementary material for "A multiscale model of complex endothelial cell dynamics in early angiogenesis": S2 Table

| Parameter | Value | Parameter | Value | Parameter | Value | Parameter | Value |
|---|---|---|---|---|---|---|---|
| $R_c$ | $1.5h$ | $D_\omega$ | 1.0 | $c_{max}$ | 1.0 | $\Delta_l$ | 0.01 |
| $E_{F1}$ | 0.25 | $E_{F2}$ | 0.7 | $s_{F1}$ | 35.0 | $s_{F2}$ | 10.0 |
| $p_{max}$ | 0.26 | $s_p$ | 0.0015 | $D_p$ | 1500 | $a$ | 7.0 |
| $n$ | 2 | $K$ | 20.0 | $k_m$ | 2.6 | $k_D$ | 0.0002 |
| $\eta_l$ | 0.1 | $\eta_{max}$ | 12.5 | $s_c$ | 0.003 | $D_c$ | 4200 |
| $\gamma_{max}$ | 17.0 | $s_m$ | 0.003 | $D_m$ | 4200 | | |

**S2 Table. Parameter values of the cellular and tissue scales used in our simulations.**
