## Supplementary material for "A multiscale model of complex endothelial cell dynamics in early angiogenesis": S3 Table

| Variable | Indexes | Description |
| --- | --- | --- |
| $E_i = 1$<br>$E_i = 0$ | $i \in \mathcal{I}_{init}$<br>$i \in \mathcal{I} \setminus \mathcal{I}_{init}$ | Initial distribution of cell nuclei. |
| $N_i = \text{Unif}[(1-\xi)b_N, (1+\xi)b_N]$<br>$D_i = \text{DUnif}[(1-\xi)b_D, (1+\xi)b_D]$<br>$I_i = \text{DUnif}[(1-\xi)I_0, (1+\xi)I_0]$<br>$R2_i = \text{DUnif}[(1-\xi)b_{R2}, (1+\xi)b_{R2}]$<br>$R2^*_i = \text{DUnif}[(1-\xi)R2^*_0, (1+\xi)R2^*_0]$<br>$N_i = D_i = I_i = R2_i = R2^*_i = 0$ | $i \in \mathcal{I}_{init}$<br>$i \in \mathcal{I} \setminus \mathcal{I}_{init}$ | Cells are initialised with ligand/receptor numbers corresponding to their baseline gene expression with a correction for random fluctuations included via the parameter $\xi$. At the voxels where there is no cell nucleus, subcellular variables are initialised with the value zero. |
| $l_i^{s_{init}} = 2\Delta_{init}$ | $i \in \mathcal{I}_{init}$ | The alignment of ECM fibrils for voxels where cells were initially placed in the direction $s_{init} \in \mathcal{S}$. |
| $l_i^s = \text{Unif}[0, \Delta_{init}]$<br>$l_i^s = \text{Unif}[0, \Delta_{init}]$ | $i \in \mathcal{I} \setminus \mathcal{I}_{init}$, $\forall s \in \mathcal{S}$<br>$i \in \mathcal{I}_{init}$, $\forall s \neq s_{init} \in \mathcal{S}$ | The alignment of ECM fibrils for the rest of the voxels is initialised with a small random value in a given range, $[0, \Delta_{init}]$, imitating random orientation of fibrils prior to their realignment due to cell migration. |
| $c_i = c_{init}$<br>$c_i = c_{max}$ | $i \in \mathcal{I}_{init}$<br>$i \in \mathcal{I} \setminus \mathcal{I}_{init}$ | The ECM concentration at the voxels with cells is equal to $c_{init} \in [0, c_{max}]$ (specified for each numerical experiment). For other voxels, the ECM is assumed to be unchanged, thus equal to the maximum ECM concentration, $c_{max}$. |
| $m_i = m_{init}$<br>$m_i = 0$ | $i \in \mathcal{I}_{init}$<br>$i \in \mathcal{I} \setminus \mathcal{I}_{init}$ | The concentration of BM components at the voxels with cells is equal to $m_{init} \in [0, 1]$ (specified for each numerical experiment). For other voxels, no BM components have been deposited, thus the concentration is set to zero. |

**S3 Table. Initial conditions for numerical simulations.** Here $\mathcal{I}$ is the set of all voxels; $\mathcal{S}$ is the set of all possible migration directions. DUnif$[a, b]$ is a discrete uniform distribution over all integer numbers lying within the interval $[a, b]$; Unif$[a, b]$ is the uniform distribution on the interval $[a, b]$. Baseline gene expression parameters for the VEGF-Delta-Notch signalling are listed in S1 Table. $\Delta_{init} = 1.0$ for all numerical simulations (this value, as, in general, for the value of the OL variable, is non-dimensional). The fluctuation parameter, $\xi$, is set to 0.1 in all numerical simulations. The exact values for $c_{init}$ and $m_{init}$ are given for each numerical experiment in S4 Table, as well as the set of initial cell positions, $\mathcal{I}_{init}$. For the description of model variables see Table 1 in the main text.
